# The Lawn is Buzzing: Increasing insect biodiversity in urban greenspaces through low-intensity mowing

**DOI:** 10.1101/2025.09.23.677970

**Authors:** Luis Mata, Drew Echberg, Charlotte Napper, Amy K. Hahs, Estibaliz Palma

**Author notes:** Corresponding author: Luis Mata (;).

## Abstract

Greenspaces have become the anvil where stewards and practitioners are forging innovative, evidence-based actions to meet biodiversity targets in urban environments, catalysing a wave of co-designed research/practice projects aimed at assessing the ecological changes brought about by urban greening and generating the evidence that biodiversity objectives are being met. Their full potential often remains unrealised due to entrenched management practices, as best exemplified by high-intensity mowing, which has given rise to the most ubiquitous greenspaces feature worldwide: the turfgrass lawn. Lawns are notoriously deficient at supporting insect communities due to their simplified vegetation structure and low plant diversity, and the compounded effect of frequent mowing on forb growth, which limits their capacity to come into flower and supply floral resources to pollinators and other flower visitors. Addressing these shortcomings can be readily achieved by reducing mowing intensity, resulting in greater vegetation height, flower cover and plant diversity – effectively transforming lawns into a more complex grassland-type ecosystem. This approach is particularly enticing to practitioners pursuing positive, cost-effective biodiversity outcomes while upholding their commitment to core ecological restoration and biodiversity conservation projects.

Here, we demonstrate how transitioning a lawn from high-to low-intensity mowing regimes led to pronounced increases in the number of indigenous insect species, evident both for the whole community and on assemblages of functionally similar species, including detritivores, herbivores, predators, parasitoids and pollinators. We further identify a positive effect of vegetation height on the community and species-specific probabilities of occurrence of indigenous species, which was consistently strong for detritivores, herbivores and parasitoids. We also show that the number of indigenous species associated with our low-intensity mowing treatment markedly exceeded that of 43 high-intensity mowed lawns previously surveyed throughout the study area, and that the effect of vegetation height across our field experiment gradient was substantially stronger than that of the existing high-intensity lawns gradient. Our findings provide compelling evidence that reducing lawn mowing intensity yields positive ecological outcomes for functionally diverse indigenous insect communities, charting a course for stakeholders tasked with demonstrating how evidence-based greening actions can be a sound investment to meet local, regional and global biodiversity targets.

## Introduction

The idea that urban greenspaces benefit both biodiversity and people has become a core tenet of urban ecological research and practice. Urban greenspaces around the world – from large remnant bushland reserves (Ofori et al. 2018) and golf courses (Threlfall et al. 2017) to small memorial sites (Mata et al. 2023) and nature strips (Bell et al. 2025), from permanent green roofs (Fenoglio et al. 2023) to temporary pop-up parks (Mata et al. 2019), and from private residential gardens (Baldock et al. 2019, Majewska and Altizer 2020) to public parks (Fan et al. 2023) and city squares (Fairbairn et al. 2024) – are providing food and habitat resources to functionally diverse microbial, fungal, plant and animal communities. Synergistically, greenspaces enrich people’s lives by providing a wide-ranging array of interwoven health, well-being, social, climate and cultural benefits (Lai et al. 2019, Mata et al. 2020, Stevenson et al. 2020, Mumaw and Mata 2022b, Li et al. 2024b, Davis et al. 2025, Kowarik et al. 2025). Not surprisingly, policymakers and leaders across all levels of grassroot and government organisations are enshrining mandates and targets aimed at protecting existing nature and/or bringing nature back into urban environments (Government of South Africa 2015, Nilon et al. 2017, United Nations 2017, World Health Organization 2017, Bush 2020, Gardens for Wildlife Victoria 2020, Pierce et al. 2020, European Union 2021, Commonwealth of Australia 2024, United States Environmental Protection Agency 2025). Greenspaces have thus become the anvil where stewards, practitioners and professionals – including community champions, school educators, friends-of-groups, volunteers, landscape architects, urban designers, city planners and sustainability consultants – are forging innovative, creative and evidence-based greening actions to operationalise ‘Nature in the City’ charters, plans, strategies and policies (Apfelbeck et al. 2020, Mata et al. 2020, Soanes et al. 2023, Kowarik et al. 2025). In line with this, greenspaces have also catalysed an emerging wave of collaborative co-designed research/practice projects focusing on assessing the ecological changes brought about by greening actions and providing the scientific evidence that specific biodiversity objectives are being met (Cadotte et al. 2017, Apfelbeck et al. 2020, Kurle et al. 2022, Brown et al. 2024, Mata et al. 2024).

The tremendous potential of greenspaces to advance the 21st-century social, biodiversity and sustainability agendas remains in many cases largely unrealised due to the legacy of pervasive and recalcitrant management practices (Aronson et al. 2017). One of the most entrenched of these is the high-intensity mowing, weeding, pest-control and watering management approach that has given rise to the most ubiquitous feature of greenspaces worldwide: the turfgrass lawn – the ultimate nemesis of a thriving urban ecosystem. Lawns are notoriously deficient at providing food and habitat resources for consumers, as well as higher trophic levels, with a substantial body of research showing that the simplified vegetation structure and low plant diversity of lawns leads to depauperated insect and other invertebrate communities, particularly when compared to alternative management approaches (Norton et al. 2019, Watson et al. 2020, Proske et al. 2022, Fekete et al. 2024) or when compared side-by-side with the insect communities supported by other vegetation layers (e.g. midstorey, tree canopy) and growth forms (e.g. tussock grass, shrubs) (Mata et al. 2021). For example, a survey conducted in the central municipality of Melbourne, Australia found lawns were associated on average with 2.5 and 4.3 times fewer indigenous insect species than midstorey and tussock grass species, respectively (Mata et al. 2021). The direct detrimental effects of the lawns’ lack of vegetation complexity and diversity on greenspace biodiversity is further compounded by the effect of mowing intensity on forb growth, which limits the capacity of forbs to come into flower and supply nectar, pollen and other floral resources to insect pollinators and other flower-visitor species (Cloutier et al. 2024, Fekete et al. 2024, Rada et al. 2024, Biella et al. 2025). Yet, high-intensity lawn management continues largely unchallenged, despite unambiguous evidence of its poor ecological performance. Maintained almost exclusively for their aesthetic and recreational values, lawns remain the defining element of urban greenspaces, covering as much as 50–70% of planted areas independent of climate, water availability and cultural contexts (Ignatieva and Hedblom 2018, Ignatieva et al. 2020).

An evolving spectrum of innovative and creative solutions have been conceptualised and put into practice around the world to address the ecological and biodiversity shortcomings of lawns (Ignatieva et al. 2020). One frequently explored pathway is to replace the lawn with an entirely new plant community, a practice that has been put into gear by scraping the lawn and either (1) sowing seed mixes – typically to achieve ecological successions leading to restored or novel grassland-, meadow- and prairie-like ecosystems (Jiang & Yuan 2017, Norton et al. 2019, Marshall et al. 2023, Horsfall et al. 2024) – or (2)planting seedlings of species that have been identified as part of a targeted design plant community – typically to achieve low-maintenance, perennial communities such as woody meadows (Walls and Greene 2023) and xeriscape, verge and pollinator gardens (Sharath and Peter 2019, Marshall et al. 2019, Mody et al. 2020, Mata et al. 2024, Bell et al. 2025). These and other related approaches (e.g. incorporating pollinator-attracting groundcovers into existing lawns; Wolfin et al. 2023, Grenier et al. 2025) epitomise innovation in biodiversity- and sustainability-oriented urban greening practice. However, their full uptake remains low due to project complexity and funding implementation barriers.

An alternative pathway that presents key economic and application advantages entails bringing vegetation complexity and plant diversity back into the lawn by reducing mowing intensity. Previous research has consistently shown that reducing mowing frequency allows the lawn to flourish in both vegetation height and flower cover, while leading to concomitant pronounced increases in plant taxonomic, functional and phylogenetic diversity – thus effectively transforming the lawn into a much more ecologically complex grassland-type ecosystem (Garbuzov et al. 2015, Rudolph et al. 2017, Chollet et al. 2018, Sehrt et al. 2020). This approach is particularly enticing to practitioners seeking to achieve positive, cost-effective biodiversity outcomes while remaining strong in their commitment to plan, fund and execute more complex and costly ecological restoration (e.g. restoration of indigenous grasslands) and biodiversity conservation (e.g. re-introduction of threatened species) projects.

It is worth emphasising that the literature widely supports the view that low-intensity mowing can lead to positive ecological outcomes. Notably, a series of recent syntheses have brought into focus the clear positive effects of reduced mowing frequencies (two or fewer times a year) on the abundance and taxa richness of insects and other arthropods (Watson et al. 2020, Proske et al. 2022, Fekete et al. 2024, Hu & Lima 2024). For instance, as much as 83% of the studies included in Fekete and colleagues’ (2024) review of studies evaluating the effects of mowing intensity on insect and other arthropod groups found that reductions of mowing frequency had a positive effect on the abundance and/or diversity of the targeted species or taxa. A more recent multi-taxon investigation by Biella and colleagues (2025) reported that areas mowed less intensively were associated on average with 1.6 times as many bee, beetle, butterfly, grasshopper, heteropteran bug and wasp species as the intensively mowed areas in their study. Similarly, two recent studies investigating the effect of mowing intensity on the species richness of insect pollinators and other flower-visitor species documented higher species richness in the less intensively mowed lawns (Cloutier et al. 2024, Rada et al. 2024). Taken together, this body of work provides a compelling initial evidence-base for practitioners seeking to implement low-intensity mowing actions aimed at supporting thriving insect and other invertebrate communities in urban greenspace lawns.

Key conceptual and methodological gaps remain to be addressed, however, to bring into full focus the ecological and biodiversity benefits of shifting lawn management paradigms to adopt and prioritise low-intensity mowing. A critical concern is the pronounced geographical bias permeating the literature, with peer-reviewed research to date primarily focused on the effects of lawn mowing intensity on non-plant biodiversity conducted in either Europe or North America. Another key issue is the wide heterogeneity displayed in terms of both survey methodologies and specimen identification. For example, some studies used passive methods (e.g. pan-trapping), which are likely to attract species from outside the survey’s sampling units (e.g. experimental or observational plots), as opposed to active methods (e.g. sweep-netting, direct observation) that fully capture the associations between the targeted taxa and the sampling units where they are being surveyed (e.g. plots, plant individuals). Of these latter studies, most reported the taxonomic diversity of the sampled individuals at coarse taxonomical levels (e.g. order), thus only allowing the effects of mowing intensity on diversity to be reported in terms of taxa-richness, as opposed to species richness. Only a few studies partially or fully identified collected specimens to species/morphospecies – either in situ or in the laboratory – yet most omitted providing the taxonomical identification level achieved and/or only used the morphospecies-level data to produce coarsely-resolved taxonomical aggregates (e.g. wild bees, flies) for their analyses. Critically, no study to date, as best as we understand, has attempted to (1) work with the full gamut of species collected/ observed through their designated survey approach, thereby enabling more holistic, community-level inferences; and (2) use fine-level taxonomical identifications to stratify species by their ecological functional roles, allowing for a more nuanced understanding of assemblage-level responses (Belitz et al. 2025). Both these advances will bring us closer to a more refined understanding of whether the species-specific and community-level responses of insects occurring in lawns are modulated by mowing intensity, and whether these responses are shared across assemblages of functionally similar species (e.g. beetle and fly detritivores, fly and wasp parasitoids). Finally, we note the missed opportunity to purposely link the experimental design with attempts to draw inferences on the ecological mechanism(s) (e.g. increased vegetation height as a proxy of increased food provisioning and habitat complexity) likely to be more closely responsible for driving the observed diversity related changes; for instance, most studies to date are based on opportunistic vegetation height treatments that were not explicitly quantified.

Here, we show that transitioning an urban greenspace lawn from high-to low-intensity mowing led to pronounced increases in insect species richness across a diverse range of taxonomical and functional groups. We further demonstrate that the insect species richness associated with this low-intensity mowing lawn markedly exceeded that of more than 40 high-intensity mowed lawns surveyed in the adjacent municipality. Our investigation is supported by two empirically derived datasets. The first was compiled specifically for this study and comprises the species-specific occupancy of over 200 detritivore, herbivore, predator, parasitoid and pollinator insect species, as recorded across two experimental lawn plots: a low-intensity mowing treatment plot and a high-intensity mowing control plot. We used this dataset to assess how (1) insect species richness varied between the treatment and control plots and (2) the effect of vegetation height on the community and species-specific probabilities of occurrence. These assessments were conducted for the whole insect community, as well as for five assemblages of functionally similar species: detritivores, herbivores, predators, parasitoids and pollinators. The second was compiled as part of a previously published study conducted in the neighbouring municipality (Mata et al. 2016, 2021), with the subsample of the data that we use here comprising the species-specific occupancy of approximately 100 insect species, as recorded in 43 high-intensity mowing lawn plots. We use this dataset to assess how (1) the insect species richness of these plots compares to our field experiment’s treatment and control plots and (2) the effect of vegetation height on the community-level probability of occurrence compared to that estimated for our field experiment. In all our assessments, we harness the power of hierarchical community models, which are uniquely suited to assess community- and assemblage-level responses by using the collective community/assemblage data to inform the probabilities of occurrence for all observed species – including those that were rarely recorded – while simultaneously accounting for imperfect detection by integrating detection-nondetection data (Zipkin et al. 2010, Kéry and Royle 2016, Devarajan et al. 2020, Belitz et al. 2025). Critically, our study was strategically co-designed between a team of local government practitioners and ecological researchers specifically to bridge the science-practice gap (Cadotte et al. 2017, Kurle et al. 2022); that is, to ensure our study was grounded in rigorous scientific theoretical and methodological foundations, while being synergistically designed to fulfil applied objectives and yielding insights that may be readily translated into actionable management innovations that can be generalised beyond the study’s context. As such, our study charts a course for researchers, practitioners and other built-environment professionals working collaboratively to demonstrate how evidence-based urban greening innovations can be a sound investment – not only in meeting local biodiversity targets, but also more overarching mandates set out in regional and global sustainability policies.

## Methods

### Field experiment: The Lawn is Buzzing project

The field experiment component of the study was conducted as part of *The Lawn is Buzzing* (henceforth TLB), a practice/research project co-designed between conservation and biodiversity officers at Merri-bek City Council and ecologists at Lookfar Research and The University of Melbourne.

### Study design

The field experiment took place across three years (2023-2025) at Gilpin Park, a public greenspace in the City of Merri-bek, Melbourne, Victoria, Australia (Appendix S1: Fig. S1). The park is approximately 65,000 m^2^ and largely covered by a high-intensity managed lawn composed of introduced grass and forb species. The park is embedded in a dense urban matrix and surrounded by lawned recreational and sports fields, mid-rise industrial facilities, and low-rise residential buildings and houses (Appendix S1: Fig. S2). There are a series of small greenspaces (2,000 – 12,000 m^2^) and sports field complexes (90,000 – 300,000 m^2^) within 200 – 1,000 m of the park, and one relatively large greenspace (1,000,000 m^2^) within 1,400 m of the park (Appendix S1: Fig. S2).

We collected insect data at two experimental lawn plots representing a low-intensity mowing (Treatment) and high-intensity mowing (Control) – of approximately 2,400 m^2^ each (Appendix S1: Fig. S3). The treatment plot was part of a network of low-intensity managed demonstration sites established across the City of Merri-bek in 2023, which were removed from their regular mowing schedule to test management implications and gather public feedback. Specifically, the treatment plot was left unmowed from October 2023 until 21 June 2024, when it was mowed to a height of about 10-15 cm, and then left unmowed for the remainder of the experiment. The control plot was established in October 2024 at the same time as the first ecological survey. It was separated from the treatment plot by 10 m (Appendix S1: Fig. S3). Across the last few years, this plot has experienced the planned, periodic mowing regime used by Merri-bek City Council in Gilpin Park. The plot continued to be mowed across the study according to the regular monthly mowing schedule.

We conducted five insect surveys at each of the experimental plots between October 2024 and April 2025. Surveys were timed as close as possible to the day(s) before the park was scheduled to be mowed, thus aiming for the lawn within the control plot to be as tall as possible within the limitations of the current mowing regime. Surveys were conducted between 10:30 and 17:30 on clear, sunny days with less than 50% cloud cover, and discontinued if rain developed or wind speed was greater than 5 m/s.

### Insect surveys

The survey protocol consisted of two components: sweep netting and direct observations. The sweep netting was aimed at documenting a wide range of insect groups known to be representative of insect communities in above ground vegetation, including ants, bees, beetles, booklice, earwigs, flies, grasshoppers, heteropteran bugs, jumping plant lice, leaf- and treehoppers, parasitoid wasps, planthoppers, and stinging wasps. We divided the experimental plots into two subplots (A and B) of equal area (200 m) and employed the same entomological net – central rod: 90 cm; bag diameter: 50 cm; bag depth: 55 cm – and method – five sweeps per each cubic metre of above-ground vegetation to achieve a survey effort proportional to the subplots’ vegetation volume – described by Mata and colleagues (2021) to collect two independent detection sub-samples during each of the five surveys. Collected specimens were transferred to labelled storing vials containing a preservative liquid (100% Ethanol). Specimens were processed in the laboratory into the project’s reference collection and identified to species/morphospecies. We then assigned these as (1)either indigenous or introduced to the study area and (2) one of the following functional groups: detritivores, herbivores, predators, and parasitoids (Appendix S1: Table S1).

Direct observations were aimed at recording the interactions (i.e. physical contact) between the plant species that were in flower during the study (Appendix S1: Table S2) and insect pollinators and other flower-visitor insects (henceforth pollinators; Appendix S1: Table S3). We used a variation of the method described by Mata and colleagues (2024) to survey up to three individuals – or group of closely co-located (within 1 m^2^) individuals – of each plant species flowering at the experimental plots, noting that within our study’s timeframe plants were only observed flowering during the two surveys conducted within the austral spring (October and November). Each individual survey consisted of three consecutive time periods of four, three, and three minutes, respectively. During each period, we actively observed the plant’s flowers, noting down the first sighting of any pollinator that came in touch with the flower’s reproductive organs (i.e. carpels and stamens). We identified the observed pollinators to species/morphospecies based on morphological features visible while they were either flying from/to the flower, stationary on the flower (i.e. basking, perching), or actively foraging the flower resources. None of the pollinators documented through this approach were, therefore, harmed during this study, noting, however, that for some taxa it was not possible to achieve species-level identification in the field – e.g. all potential sweat bee and hoverfly species were recorded as genus *Lasioglossum* and family Syrphidae, respectively (Appendix S1: Table S3).

### Photographic collection and iNaturalist community identification

To complement the field and laboratory identifications, we photographed observed interactions and collected specimens using a digital single-lens reflex camera equipped with a 100 mm macro lens. While we aimed to photograph all observed interactions and collected specimens this was not always possible, as some pollinators flew away before they could be photographed and some small specimens in the reference collection were beyond the magnification capacity of our equipment. We then uploaded the images to iNaturalist and used the identifications provided by platform’s Community Taxon system (Palma et al. 2024) to either improve or validate the taxonomic resolution of our own identifications.

### Vegetation height

To derive the vegetation height covariate, we used a metric tape measure to take 20 height measurements in each experimental plot (ten in each subplot) during each survey. These were then averaged to obtain a single vegetation height covariate for each plot (treatment/control) and survey (1:5) combination.

### Comparative lawn dataset: The Little Things that Run the City project

The comparative lawn dataset used in this study was sourced from *The Little Things that Run the City* (henceforth TLT), a research and outreach project focused on the ecology, biodiversity and conservation of insects in the City of Melbourne, which is a municipality located directly south of the City of Merri-bek (Appendix S1: Fig. S2). The subsample of the dataset we use here includes insect data collected with the same sweep netting methodological approach used in the field experimental component of this study. The data were collected across three surveys conducted between January and March 2015 in 43 lawn plots located within 15 public parks (Mata et al. 2016, 2021; Appendix S1: Table S4, Fig. S4). The subsampled dataset also includes the five vegetation height measurements that were taken in each plot during each survey, which were averaged to obtain a single vegetation height covariate for each plot and survey combination.

### Data analysis

#### Modelling species richness and the effect of vegetation height

To assess how insect species richness varied between the treatment and control plots (TLB/TLT sweep netting and TLB direct observation data) and the effect of vegetation height on the community and species-specific probabilities of occurrence (TLB/TLT sweep netting data), we used variations of the hierarchical metacommunity models (Kéry and Royle 2016) described by Mata and colleagues (2021, 2024). For the modelling of the species collected through sweep netting in the TLB project, the two plot (treatment/ control) times five survey combinations constituted the unit of analysis for drawing inferences on species occupancy and the subplots (A and B) constituted the unit of detection replication. For the modelling of the species collected through sweep netting in the TLT project, the 43 plots times three survey combinations constituted the unit of analysis for drawing inferences on species occupancy and the number of surveyed subplots within each park – which varied between 1 and 9 depending on the park area (Mata et al. 2021) – constituted the unit of detection replication. For the modelling of the species collected through direct observation in the TLB project, the two plot (treatment/control) times two surveys times nine plant species constituted the unit of analysis for drawing inferences on species occupancy and the three survey periods times the number of plant individuals surveyed – which varied between 1 and 4 depending on the plant species – constituted the unit of detection replication. Both models are organised in three levels: a first level for species occupancy; a second for species detectability; and a third to treat the occupancy and detection of each species as random effects (Kéry and Royle 2016).

The occupancy (Z) and detection (Y) levels were specified as:

Z_*i,j*_ ∼ Bernoulli (Ψ_*i,j*_)

Y_*i,j,k*_ ∼ Bernoulli (p_*i,j,k*_ · Z_*i,j*_)

where Ψ_*i,j*_ is the probability that species *i* occurs at inference point *j* and p_*i,j,k*_ the probability of species *i* being detected at inference point *j* at/during detection replicate *k*.

The linear predictors were specified on the logit-probability scale as:

logit (Ψ_*i,j*_) = occ_*i*_ + eff_*i*_ · vh_*j*_ [TLB/TLT sweep netting data]

logit (Ψ_*i,j*_) = occ_*i*_ [TLB direct observation data]

logit (p_*i,j,k*_) = det_*i*_ [TLB/TLT sweep netting and TLB direct observation data]

where occ_*i*_, det_*i*_ and eff_*i*_ are the species-specific random effects, specified as:

occ_*i*_ ∼ Normal (mu.occ, tau.occ) [TLB/TLT sweep netting and TLB direct observation data]

det_*i*_ ∼ Normal (mu.det, tau.det) [TLB/TLT sweep netting and TLB direct observation data]

eff_*i*_ ∼ Normal (mu.eff, tau.eff) [TLB/TLT sweep netting data]

The metacommunity mean occupancy (mu.occ) and detection (mu.det) hyperpriors were specified as logit(Uniform (0, 1)), the metacommunity mean effect hyperprior (mu.eff) as Normal (0, 0.001), and the metacommunity precision occupancy (tau.occ), detection (tau.det) and effect (tau.eff) hyperpriors as Gamma (0.1, 0.1).

We used the latent occurrence matrix Z_*i,j*_ to estimate the species richness associated with each plot (TLB/TLT sweep netting data) or plant species (TLB direct observation data) inference point SRj through the summation:

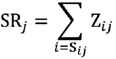

where SR_*j*_ is an indexing vector that accounts for plot-species, survey-species or plant-species specificities, that is, for each inference point, the recorded species are included with probability of occurrence = 1 and a limited random subsample of other species occurring in the study area (i.e. the other inference points) are included with their 0 < Z < 1 estimated probabilities of occurrence (Mata et al. 2021, 2024). This specification allows us to operate under the forbidden links ecological framework, where non-occurrence of pairwise interactions can be accounted for by spatiotemporal uncoupling, size and reward mismatches, and foraging, physiological and biochemical constraints (Olesen et al. 2010, Jordano 2016).

As these calculations were conducted within a Bayesian inference framework, the species by plot-survey (TLB/TLT sweep netting data) and plot-survey-plant species (TLB direct observation data) estimates were derived with their full associated uncertainties. We then averaged the richness estimates corresponding to the treatment and control inference points to obtain posterior distributions for each group that could be statistically compared.

Finally, we used the estimated community-level occupancy and effect parameters from the TLB/TLT sweep netting data models to draw predictions across a reasonable range of the vegetation height gradients (TLB: 0 – 50 cm; TLT: 0 – 8 cm).

#### Bayesian inference implementation

We applied a Bayesian inference approach to estimate model parameters, using Markov Chain Monte Carlo (MCMC) simulations to draw samples from the parameters’ posterior distributions. Models were implemented in JAGS (Plummer 2003), as accessed through the R package jagsUI (Kellner 2024; R Core Team 2024). We used three chains of 10,000 iterations, discarding the first 1,000 in each chain as burn-in. We assessed acceptable convergence by visually inspected the MCMC chains and verifying that the values of the Gelman-Rubin statistic fell below 1.1 (Gelman and Hill 2007).

## Results

### The Lawn is Buzzing: Sweep netting surveys

We recorded 194 insect species through the sweep netting surveys, including: ants (8), bees (5), beetles (32), booklice (1), brachyceran flies (44), earwigs (1), grasshoppers, crickets and katydids (6), heteropteran bugs (17), jumping plant lice (3), lacewings (1), leafhoppers (16), nematoceran flies (1), parasitoid wasps (48), planthoppers (4) and stinging wasps (7) (Appendix S1: Table S1). These represented 49 detritivore, 78 herbivore, 12 predator and 57 parasitoid species (Fig. 1; Appendix S1: Table S1). As much as 97% of the recorded species were locally indigenous to the study area, with only six introduced species recorded: the spotted amber ladybird *Hippodamia variegata*, a carpet beetle in genus *Anthrenus*, a broad-nosed weevil (*Sitona discoideus*), the European earwig *Forficula dentata*, the European firebug *Pyrrhocoris apterus*, and the European honeybee *Apis mellifera* (Hymenoptera: Apidae) (Appendix S1: Table S1). The most commonly occurring species was the lawn fly *Hydrellia tritici* (Diptera: Ephydridae), accounting for 4.3% of all records.

**Figure 1.**
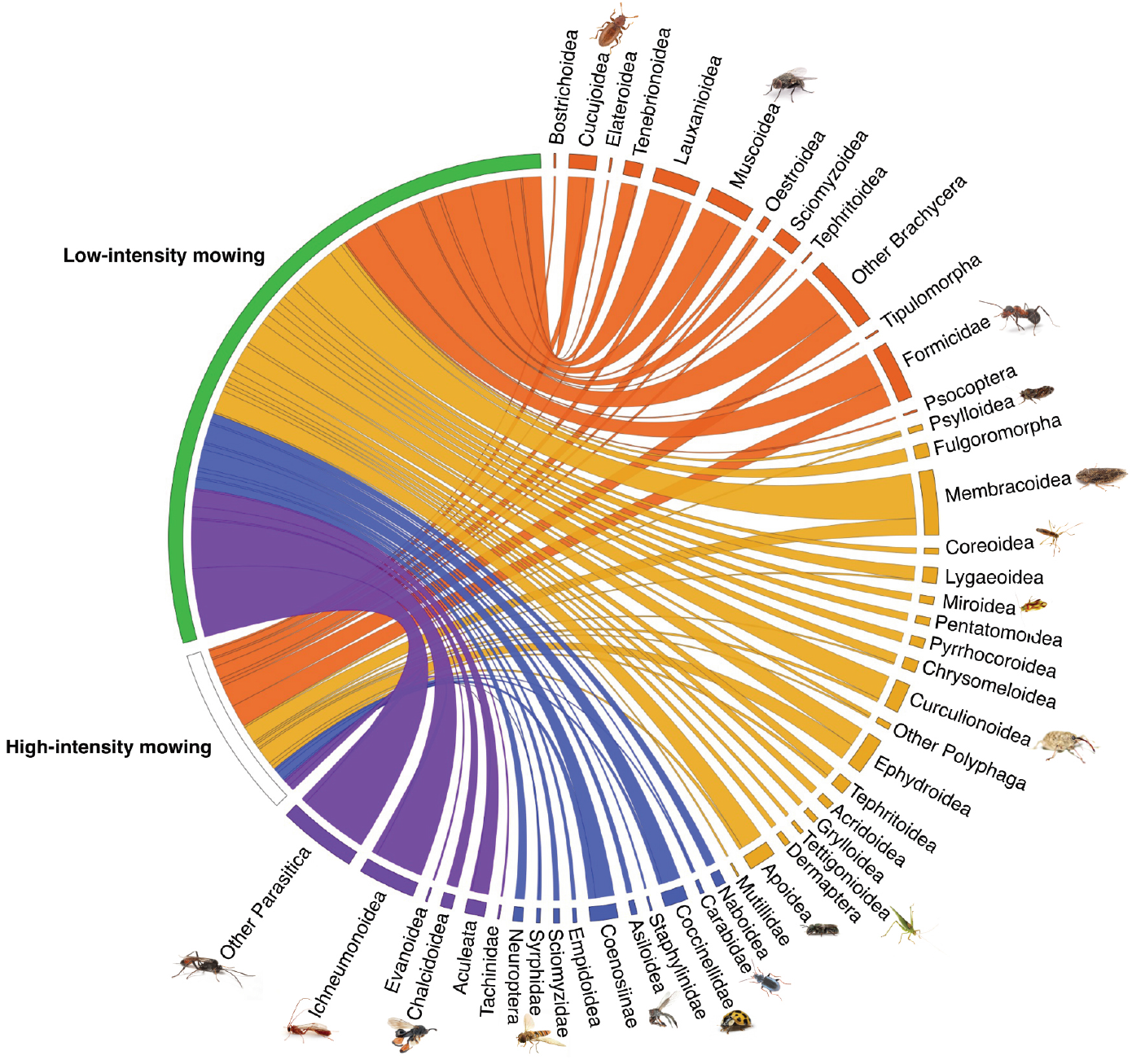
Quantitative network representation of the observed interactions between experimental lawn plots (high-intensity mowing: white box; low-intensity mowing: green box) and insect species collected through sweep-netting. Species have been organised into evolutionary clades and colour-coded into functionally similar assemblages (detritivores: orange; herbivores: yellow; predators: blue; parasitoids: purple). Chord width reflects the relative frequencies by which the insect species from the given clade are represented in the interaction.

### The Lawn is Buzzing: Direct observation surveys

We recorded 14 pollinator and other flower-visitor insect species through the direct observation surveys, including: ants and bullants (2); banded beeflies (1) fruitflies, hoverflies and other brachyceran flies (3); sweat bees and honeybees (2); butterflies (3); heteropteran bugs (2); and parasitoid wasps (1) (Appendix S1: Fig. S5, Table S3). As much as 86% of the recorded species were locally indigenous to the study area, with only two introduced species recorded: the spotted amber ladybird *H. variegata* and the European honeybee *A. mellifera* (Appendix S1: Table S3). The most commonly occurring pollinator were hoverflies (Diptera: Syrphidae) accounting for 29% of all records.

### The Lawn is Buzzing: Species richness

The estimated number of indigenous insect species found through the sweep netting surveys on the low-intensity mowing treatment plot was on average 5.1 times higher than that for the high-intensity mowing control plot (Fig. 2a; Appendix S1: Table S5). The marked statistical difference we found in the estimated number of species between the treatment and control plots were consistent across the four studied functional groups: detritivores (2.5 times higher), herbivores (4.3 times higher), predators (1.6 times higher) and parasitoids (6.8 times higher) (Fig 2b; Appendix S1: Table S5).

**Figure 2.**
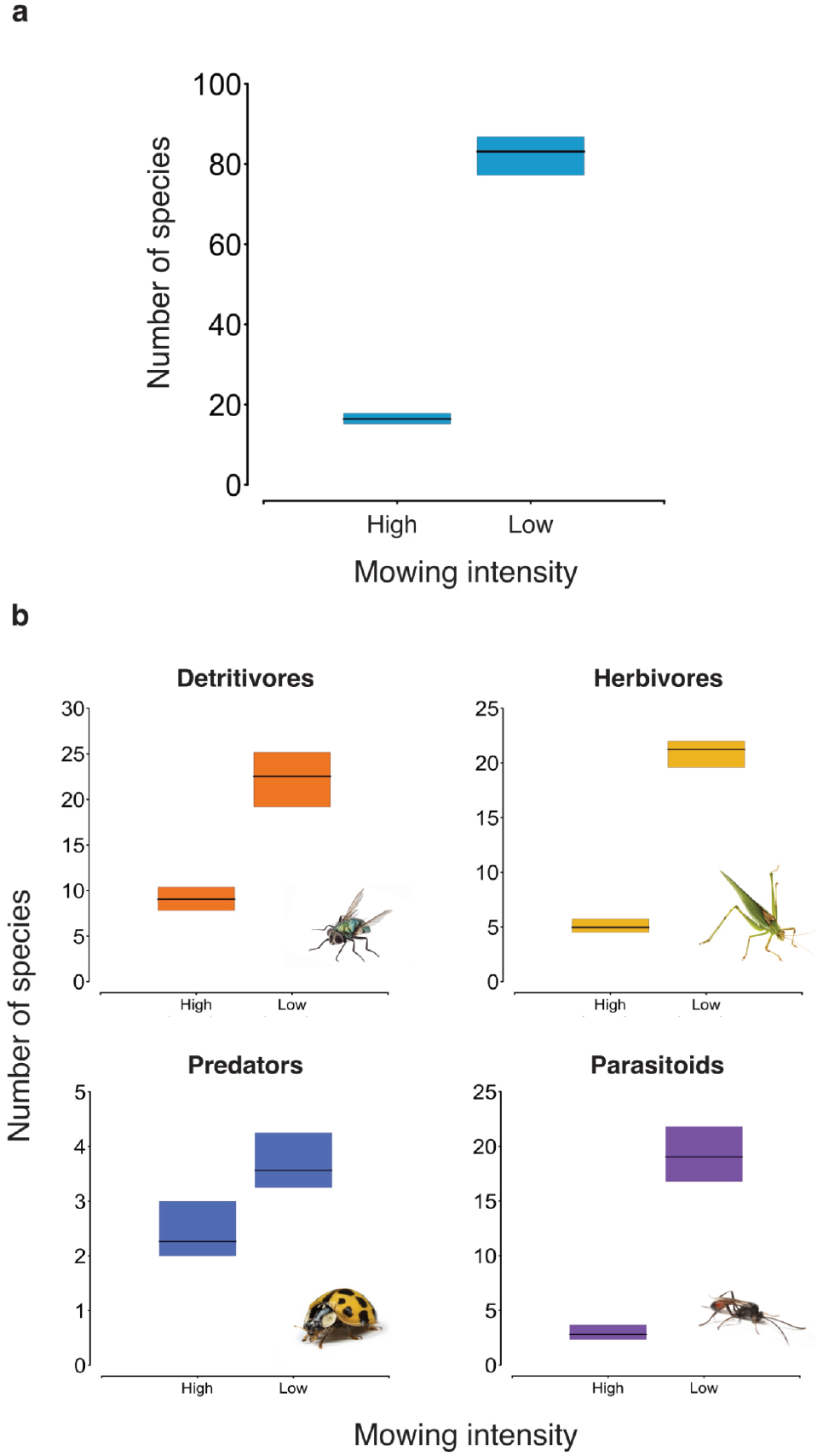
Number of indigenous insect species by mowing intensity treatment as estimated for the whole insect community (**a**) and each assemblage of functionally similar species (**b**). Black horizontal lines represent the mean number of species and the coloured vertical bars the associated uncertainty (95% credible intervals).

The estimated number of indigenous pollinator species found through the direct observation surveys on the low-intensity mowing treatment plot plant species was on average 3.5 times higher than that for the high-intensity mowing control plot plant species (Fig. 3a; Appendix S1: Table S5). In terms of individual plant species, the common dandelion *Taraxacum officinale* and the coastal galenia *Aizoon pubescens* were the plant species with the highest associated number of indigenous pollinators (Fig. 3b; Appendix S1: Table S2).

**Figure 3.**
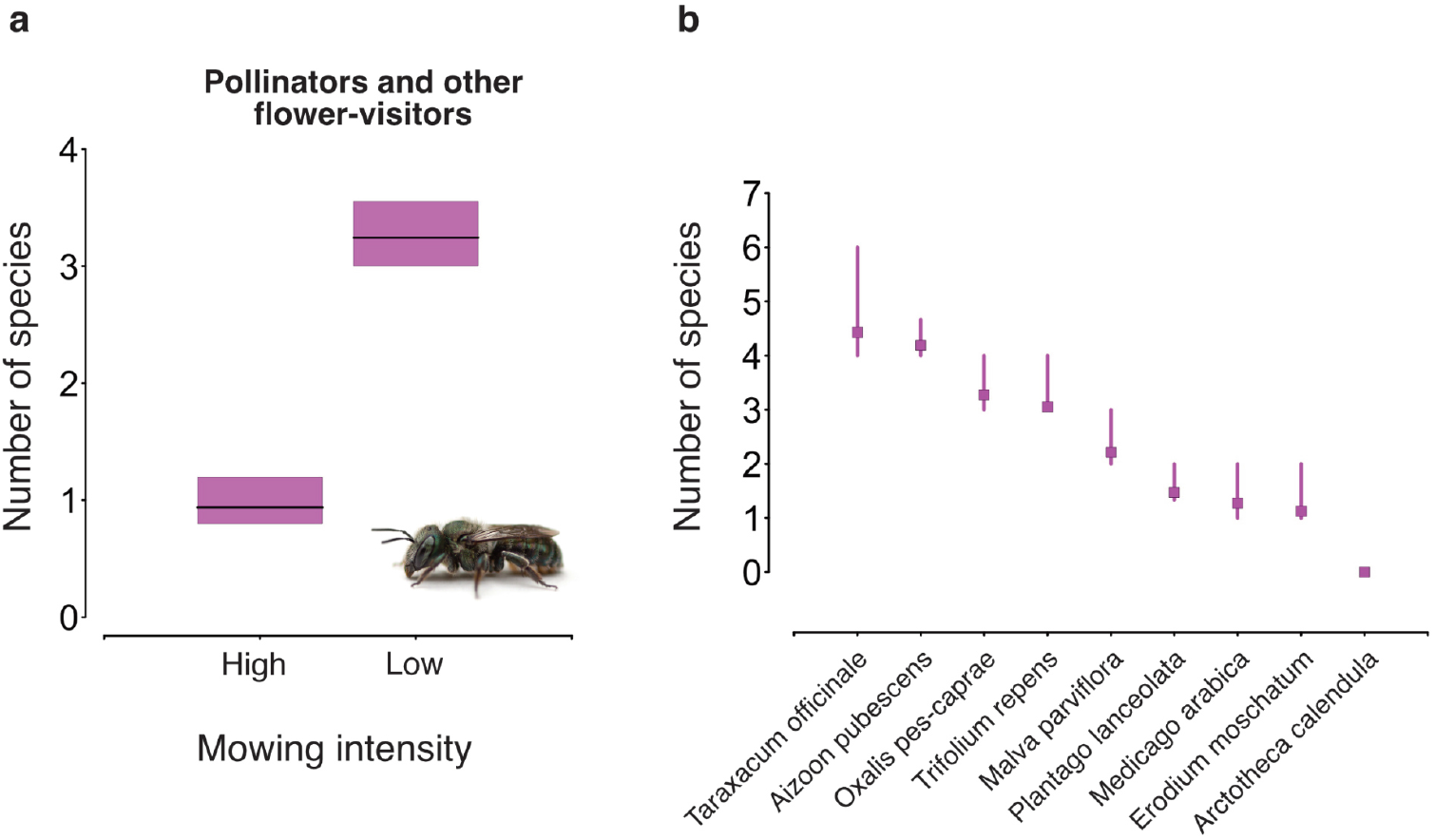
Estimated number of indigenous pollinators and other flower-visitor insect species by (**a**) mowing intensity treatment and (**b**) by plant species. The black horizontal lines (**a**) and the coloured squares (**b**) represent the mean number of species; the coloured vertical bars (**a**) and vertical lines (**b**) represent the associated uncertainty (95% credible intervals).

### The Lawn is Buzzing

Effect of vegetation height Vegetation height had a strong positive effect on the community-level probability of occurrence of indigenous species (Fig. 4a; Appendix S1: Table S6), with as many as 94% of the species showing positive effects with 95% Credible Intervals that did not overlap zero (Appendix S1: Table S7). This positive effect of vegetation height on the community-level probability of occurrence of indigenous species was consistent for detritivores, herbivores and parasitoids, whereas the effect on predators was marginally positive with a 95% Credible Interval that overlapped zero (Fig. 4a; Appendix S1: Table S6).

**Figure 4.**
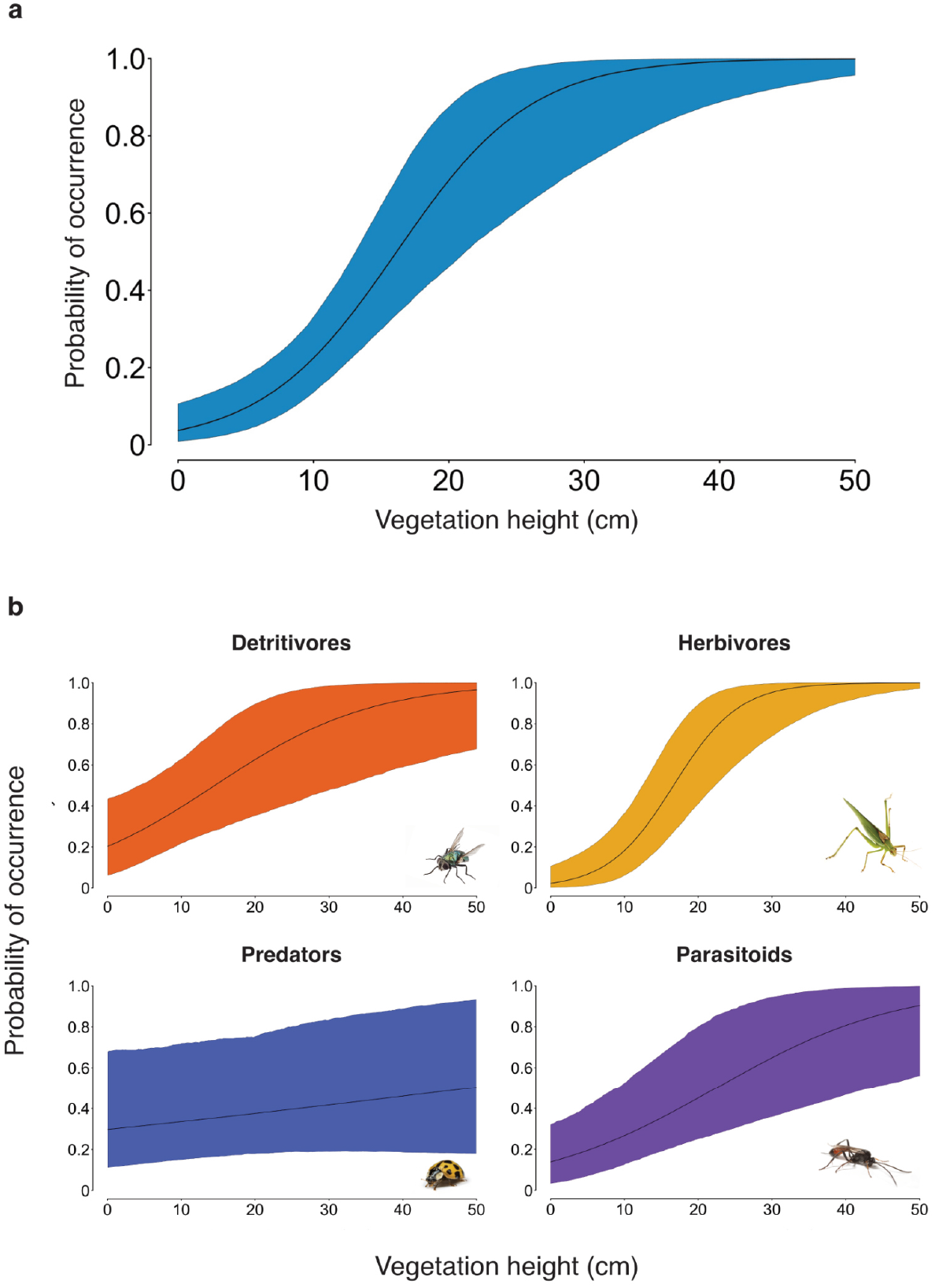
Effect of vegetation height on the community-level probability of occurrence of indigenous insect species as estimated for the whole insect community (**a**) and each assemblage of functionally similar species (**b**). Black curved lines represent the mean response and the coloured curved polygons the associated uncertainty (95% credible intervals).

#### Comparisons between The Lawn is Buzzing and The Little Things that Run the City

When the species richness estimates for the lawn plots from both *The Lawn is Buzzing* and *The Little Things that Run the City* projects were compared side-by-side, it became evident that the low-intensity mowing treatment plot from the TLB project was associated with the highest number of indigenous insect species (Fig. 5a; Appendix S1: Table S8). This plot showed a species richness that was on average 2.3 times higher than the plot from the TLT project associated with the highest number of species (Fig. 5a; Appendix S1: Table S8). On the other hand, the estimated number of species in the high-intensity mowing control plot from TLB fell within the species richness range observed across the TLT plots (Fig. 5a; Appendix S1: Table S8). This plot showed a species richness that was on average higher than the species richness observed in 80% of the TLT plots (Fig. 5a; Appendix S1: Table S8).

**Figure 5.**
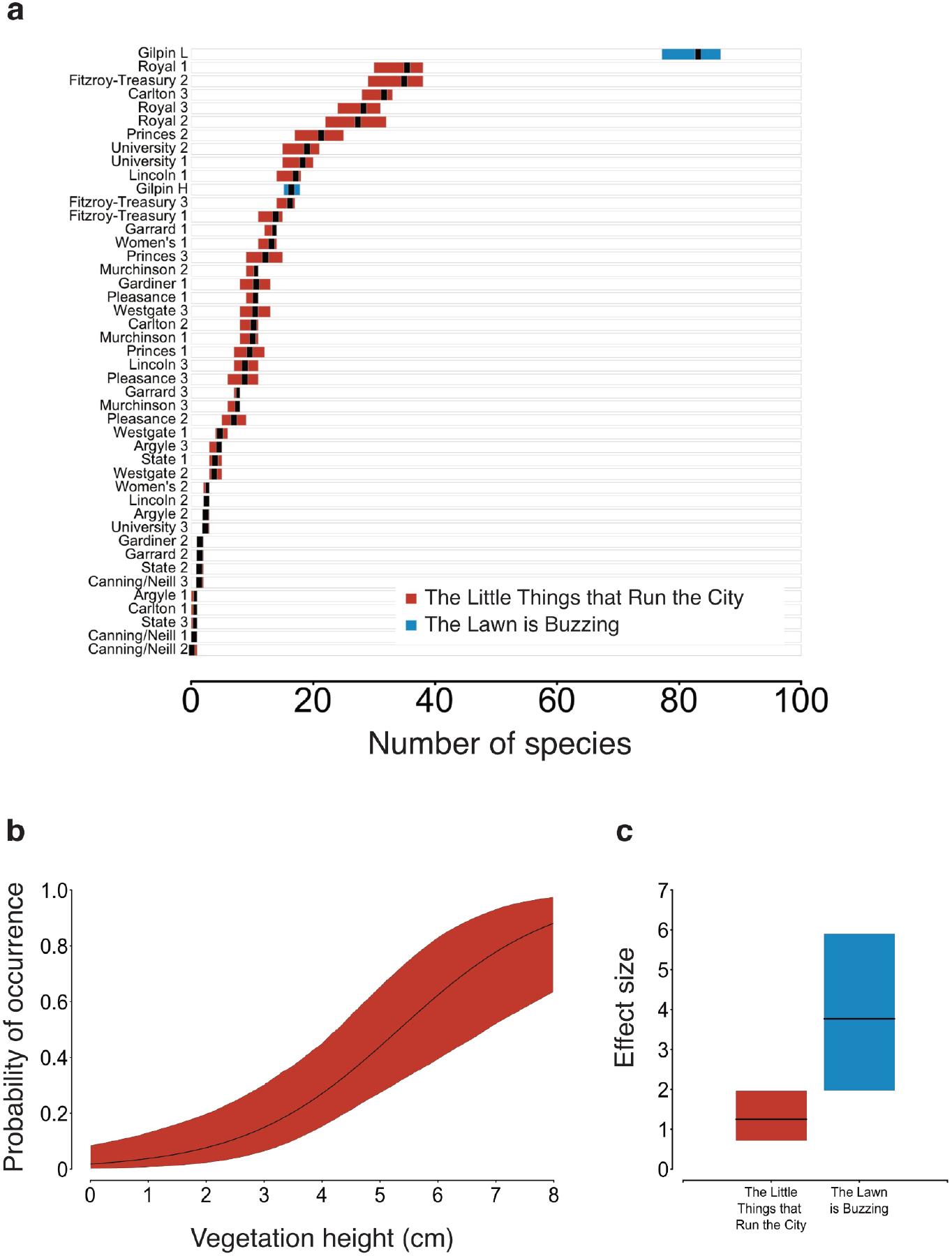
(**a**) Number of indigenous insect species by lawn plot as estimated for the 43 high-intensity lawn plots investigated in *The Little Things that Run the City* project (red) and the high- and low-intensity mowing lawn plots investigated in *The Lawn is Buzzing* project (blue). (**b**) Effect of vegetation height on the probability of occurrence of indigenous insect species as estimated for the whole insect community investigated in *The Little Things that Run the City* project. **(c**) Effect of vegetation height on the community-level probability of occurrence of indigenous insect species as estimated across the high-intensity mowing plots in T*he Little Things that Run the City* project (red) and the low- and high-intensity mowing plots in *The Lawn is Buzzing* project (blue). Black lines (**a, b, c**) represent mean responses; coloured boxes (a, c) and curved polygon (b) represent the associated uncertainty (95% credible intervals).

As in the two experimental plots in the TLB project, vegetation height across the 43 lawn plots in the TLT project also had a positive effect on the community-level probability of occurrence of indigenous insect species (Fig. 5b; Appendix S1: Table S6). When the effect of vegetation height on the community-level probability of occurrence of indigenous insect species was compared between TLT and TLB, the latter showed an effect that was on average 3.0 times higher (Fig. 5c; Appendix S1: Table S6).

## Discussion

In this study – co-designed between local government practitioners and ecological researchers – we demonstrate how transitioning an urban greenspace lawn from high-to low-intensity mowing led to a pronounced increase in the number of locally indigenous insect species. Critically, we show that the observed rise in species richness is not only seen at the level of the overall insect community, but also on assemblages of functionally similar species, including detritivores, herbivores, predators, parasitoids and pollinators. Our study further highlights the strong positive effect that our investigated causal mechanism – vegetation height as a proxy of increased food provisioning and habitat complexity – had on the community and species-specific probabilities of occurrence of indigenous insect species, an effect that was consistently strong for detritivores, herbivores and parasitoids, but only marginally positive for predators. Additionally, we show that the number of indigenous insect species associated with our low-intensity mowing experimental lawn markedly exceeded that of 43 high-intensity mowed lawns surveyed throughout the broader, adjacent urban area, and that the effect of vegetation height across the gradient investigated in our field experiment was substantially stronger than that observed across the gradient of the 43 high-intensity mowing lawns. When considered together, our findings provide compelling evidence that reducing lawn mowing intensity in urban greenspaces can produce positive ecological outcomes for indigenous insect species across a diverse range of taxonomical and functional groups.

### Low-intensity mowing increases indigenous insect biodiversity

Our study has yielded compelling evidence of the positive effect that reducing lawn mowing intensity has on supporting a diverse community of indigenous insects. Specifically, data from our low- and high-intensity mowing experimental plots show that a 6-fold reduction in lawn mowing frequency (i.e. from 12 to 2 times per year) led to an approximately 5-fold increase in the number of indigenous insect species. This finding is consistent with those reported in the recent reviews by Fekete and colleagues (2024) and Hu and Lima (2024), as well as with the quantitative syntheses by Watson and colleagues (2020) and Proske and colleagues (2022), all of which have highlighted the positive effects of reduced lawn mowing on insect species richness. However, the average positive effect size (5.08) on insect species richness we report here for low-intensity mowing is considerably larger than those reported in the latter two meta-analyses: 1.05 in Watson and colleagues (2020) and 1.25 in Proske and colleagues (2022) – noting that (1) all of the studies included in these meta-analyses were conducted in the Northern Hemisphere; (2) some of their summary effect sizes included abundance as opposed to strictly richness study-level estimates; and (3) most of their summary effect sizes were based largely on studies reporting taxa (as opposed to species) richness estimates and/or reporting species richness of a particular arthropod/ insect group (as opposed to the whole insect/ arthropod community). The more recent study conducted in Europe by Biella and colleagues (2025) – which was more closely aligned with ours in terms of its attempt to investigate the whole insect community, albeit solely relying on visual in situ morphospecies identification without any collected specimens – also found a positive effect of reduced mowing, reporting an effect size (1.6) that was consistent with the meta-analyses but again considerably lower than the one we report here. When considered together, our findings and those previously reported in the literature offer robust evidence that reducing lawn mowing intensity can contribute to support insect communities in urban greenspaces beyond what is currently achieved under high-intensity mowing. They also provide initial but compelling empirical support for this phenomenon occurring outside the North American and European contexts where it has traditionally been studied.

It should be emphasised that the positive effects of reduced lawn mowing we found for the whole insect community were consistent across assemblages of functionally similar species, including detritivores, herbivores, predators and parasitoids. This finding agrees with previous research demonstrating the capacity of urban greenspaces to support functionally diverse communities of insects and other arthropods when properly managed (Braaker et al. 2017, Mata et al. 2017, 2021, 2023, Gonzalez-Cesped 2021, Damptey et al. 2022). Critically, this multitrophic richness plays a fundamental role in driving energy fluxes through and beyond the lawn food web (Barnes et al. 2018), thus boosting the overall ecosystem multifunctionality of its associated greenspace (Lefcheck et al. 2015, Soliveres et al. 2016, Li et al. 2024a). For instance, multitrophic diverse lawn insect communities can boost plant productivity and belowground biochemical processes by cycling organic matter back into the soil, foster plant health across the entire greenspace by suppressing potential pests, and support higher-level taxa by acting as prey to both other insects (e.g. the immature stages of fly and wasp parasitoids) and a range of non-insect taxa living and/or foraging in urban greenspaces such as reptiles, birds and bats. Especially noteworthy was the strong positive effect of reduced mowing on parasitoids – an assemblage comprised largely of chalcidoid, ichneumonid and other parasitoid wasps but also including one species of tachinid fly – which showed an approximate 7-fold increase in species richness in the low-intensity mowing plot. Supporting existing and attracting new parasitoid species to urban greenspaces is particularly challenging given the more complex ecological conditions required by most species to complete their life cycles; for instance, adult parasitoid females require habitats that synergistically meet their reproductive (i.e. hosts for oviposition) and nutritional (i.e. floral resources to maximise host-finding efficacy) needs (Lewis et al. 1998). This finding therefore provides additional evidence in favour of our argument for the capacity of lawns to sustain functionally complex insect communities when released from the frequent disturbances they experience under frequent mowing management regimes.

### Low-intensity mowing attracts indigenous insect pollinators

In our assessment of insect pollinators – bees, butterflies and hoverflies, but also other flower-visitor species such as ants, ladybugs, firebugs and parasitoid wasps – we also found strong evidence of the positive effects of reduced lawn mowing. Notably, the 6-fold reduction in mowing frequency experienced by the low-mowing treatment plot led to an average 3.5-fold increase in the number of indigenous pollinator species observed on the low-compared to the high-intensity plot. While this finding largely concurs with previous studies (Cloutier et al. 2024, Fekete et al. 2024, Rada et al. 2024), our study goes a step further by demonstrating for the first time that the positive effect of reduced lawn mowing on pollinator insect richness is also realised in urban environments other than in North America and Europe in a system where the floral resources are provided exclusively by introduced forb species. Importantly, this was shown using a methodological approach that targeted the whole community of indigenous pollinator and other flower-visitor insects, rather than focusing on a single taxon or group of taxa in isolation. Our results are also consistent with previous studies demonstrating that insect pollinators benefit from restoring lawns into grassland-, meadow- and prairie-like ecosystems (Ulrich & Sargent 2025), fully replacing them with designed perennial plant communities (Brown et al. 2024, Mata et al. 2024, Bell et al. 2025), or complementing them with pollinator-attracting groundcovers (Wolfin et al. 2023, Grenier et al. 2025). Jointly interpreted, our findings and those established in prior research bring into focus the biodiversity gains for insect pollinators and other flower-visitor insects that may be reached through greening actions aimed at transitioning lawns into less intensively managed, restored, or purposefully designed habitats rich in floral resources.

Beyond the positive effects of reduced mowing, our data indicates that the number of indigenous pollinators observed on the introduced forbs that flowered during the field experiment was relatively low and varied discernibly amongst species. For instance, the common dandelion *Taraxacum officinale* and the coastal galenia *Aizoon pubescens* were associated with the highest number of indigenous pollinators (4.4 and 4.2 species on average, respectively). This means that, on average, they were associated with approximately 2.8 – 4.0 times more indigenous pollinator species than the ribwort plantain *Plantago lanceolata*, the spotted medick *Medicago arabica*, and the musk stork’s-bill *Erodium moschatum*, which were associated with the lowest number of species (1.5, 1.3 and 1.1 species on average, respectively). Notably, all the introduced forbs in our study were associated with fewer indigenous pollinator species on average than the highest-performing indigenous plant species investigated in Mumaw and Mata (2022a) and Mata and colleagues (2024) – recognising that the close similarities in geographical, landscape and methodological contexts between these two studies and ours allow for direct and robust comparisons. For example, the sticky everlasting *Xerochrysum viscosum* – the top-performing indigenous forb in both studies – was associated with 2.1 times more indigenous pollinators in Mata and colleagues (2024) and 5.1 times more in Mumaw and Mata (2022a) than those associated with *T. officinale* in our study. Interestingly, however, *T. officinale* and *A. pubescens*, were associated with: (1) a similar average number of indigenous pollinator species as a range of the forbs locally indigenous to Melbourne, as in reported in Mumaw and Mata (2022a) and Mata and colleagues (2024); and (2) a higher average number of indigenous pollinator species than almost all the Australian native forbs investigated in Mata and colleagues (2024). As such, our findings appear to support theoretical frameworks bringing attention to the potential benefits of introduced plants as novel resources for indigenous insects (Valentine et al. 2020), as well as an array of empirical studies providing evidence of the positive role that lawn-flowering forbs – including most of the species that flowered in the low-intensity mowing plot such as T. officinale and P. lanceolata – can play in providing floral resources that support indigenous and introduced insect pollinators both in their introduced (e.g. European forbs in North America: Larson et al. 2014, Lowenstein et al. 2019, Wolfin et al. 2023, Jaiswal and Joseph 2024) and native (Tew et al. 2021) ranges. The provision of floral resources such as pollen and nectar by introduced forbs may mediate positive effects beyond the lawn; for instance, the soursob *Oxalis pes-caprae* – a South African native that flowered exclusively in the low-intensity mowing plot – has been documented providing facilitative effects on the reproductive success of co-flowering indigenous plants in Portugal (Ferrero et al. 2013). We should sound a note of caution with regards of these findings, as both theoretical (Johnson et al. 2017) and empirical (Iskander 2021) studies have also drawn attention to the negative impacts on indigenous plants mediated by the floral resources introduced forbs provide to insect pollinators.

### Positive effects of vegetation height on insect occupancy

Our study provides considerable insights into the positive effects that vegetation height had on the community and species-specific probabilities of occurrence of indigenous insect species. Although not in itself a causal mechanism, vegetation height serves as a reliable and readily quantifiable proxy for the underlying ecological factors – namely, increased food availability and habitat complexity arising from reduced lawn mowing – that plausibly contributed to the observed increases in indigenous insect species richness. Specifically, vegetation height captured multiple axes of ecological variation in the structural, compositional and functional attributes of our lawn system. For example, the increased lawn heights achieved through our experimental reduction in mowing intensity likely increased the amount of plant biomass available to herbivores (e.g. weevils, leafhoppers and jumping plant lice), which likely determined the amount of animal biomass available to predators (e.g. ladybugs, damselbugs and lacewings) and, ultimately, the amount of organic material available to detritivores (e.g. ants and ant-like flower beetles). The diversity and composition of plant growth forms (e.g. grasses, forbs) reaching reproductive maturity also changed under low-intensity mowing, which altered the quantity and diversity of floral (e.g. pollen and nectar) and seed resources available to pollinators (e.g. bees and butterflies) and other flower-visitor species (e.g. parasitoid wasps), as well as granivores (e.g. seed bugs). Additionally, the low-intensity mowing increased the heterogeneity of microhabitats, which likely determined the amount and quality of shelter areas, foraging opportunities, and reproductive sites available to the lawn insect community; as well as the abundance and diversity of host species available to parasitoids (e.g. tachinid flies and ichneumonid wasps) to complete their life cycles.

Not unexpectedly, the strong positive effect of vegetation height on the community-level probability of occurrence of indigenous insect species was consistent for detritivores, herbivores and parasitoids, but, intriguingly, not for predators. While our study only captured the lawn vegetation height gradient in two management categories (i.e. high-vs low-intensity mowing), and did not explicitly attempt to experimentally assess whether a minimum vegetation height threshold or range exists at which the levels of insect species richness observed may be already realised, we note with interest that as much as 77% of the species recorded in our study occurred exclusively in the low-intensity mowing plot. Indeed, a few herbivorous (e.g. grasshoppers, heteropteran bugs, jumping plant lice and planthoppers) and most predatory (e.g. ladybugs, rove beetles, stiletto flies, robber flies, damselbugs and lacewings) and parasitoid (e.g. tachinid flies and parasitoid wasps) groups occurred exclusively or almost exclusively on the low-intensity mowing plot. For herbivores, the explanation is likely associated with access to a more diverse array of resources and microhabitats (e.g. seeds, hiding grounds from bird predators, foliage architecture conducive to camouflage) that became available only after the cessation of disturbances from high-intensity mowing, with predators and parasitoids possibly tracking the new ecological niches emerging from the resulting increase in insect herbivore diversity.

### Performance of our study’s low-intensity mowing plot beyond the field experiment

While our study’s field experiment data was modelled with appropriately replicated within-plot data – yielding robust statistical inferences that, importantly, accounted for the imperfect detection of species – the replication at the greenspace level was restricted to a single site. Until our primary data are synthesised into future reviews and/or meta-analyses, our ability to robustly extrapolate findings beyond the study’s geographical and climatic envelope remains limited, as does our confidence in determining whether the strong positive effects of reduced mowing on indigenous insect species will be consistently observed at other sites, even within or in proximity to the study area. Yet, one of the most innovative aspects of our study was the integration of our experimental primary findings with those from *The Little Things That Run the City* project (TLT; Mata et al. 2016, 2021), an earlier observational study conducted in the same broader area and using the same methodological approach. This allowed us to place and compare data from our low- and high-intensity mowing experimental plots with that derived from a robustly replicated network of high-intensity mowed lawns representing a richly diverse gradient of greenspace sizes within the same landscape context. In doing so, we achieved a more nuanced and synthetic understanding of our findings, while contributing to overcome some of the challenges intrinsic to single-site studies where broader, multiple-site replication is constrained by practical, financial and public perception barriers. Accordingly, our study’s experimental design and modelling methodology are well aligned with calls and guidelines advocating for meta-analytical thinking, supporting the synthesis of more comprehensive and generalisable evidence on the relationships between the studied practices and observed outcomes (Gerstner et al. 2017, Gurevitch et al. 2018, Ockendon et al. 2021).

Leveraging this synthesis and comparative framework, our analyses uncovered several noteworthy results. For instance, the number of indigenous insect species associated with our low-intensity mowing plot markedly exceeded that of the 43 high-intensity mowed lawns documented in TLT dataset. Indeed, the species richness of the low-intensity mowing plot was on average 2.3 times higher than the highest number of species recorded in TLT’s mown lawn plots. It is worth emphasising that a few TLT high-intensity mowing plots showed species richness estimates that were on average higher and fully statistically different than the species richness associated with our experimental high-intensity mowing control plot. This included, for example, all three plots from Royal Park, a greenspace approximately 20 times larger – and comprising a much richer range of habitats – than the site (Gilpin Park) where our current experimental plots were located. If the 5-fold increase in indigenous insect species richness that we observed in Gilpin Park between the high- and low-intensity mowing plots were to be consistent across Royal Park, one could predict that plots within this greenspace experiencing similar reductions in mowing intensity would achieve parallel positive shifts, potentially supporting average insect richness levels exceeding 136 species. Additionally, this approach also revealed that the effect of vegetation height on indigenous species occupancy across the gradient investigated in our current field experiment – where vegetation reached heights of up to 80 cm – was substantially stronger (on average 3.0 times higher) than the effect observed across the gradient defined by the 43 TLT high-intensity lawns – where vegetation only reached heights of up to 15 cm. Beyond increasing fundamental knowledge on how insect species respond to expanded ecological gradients, this result provides further support to our earlier argument on how the structural, compositional and functional changes to lawns spared from frequent mowing positively affect insect occupancy, particularly of species with more specialised niches or tracking specific food and habitat resources. Again, we predict that this effect may be at least equally sustained or potentially amplified in sites with similar or wider vegetation height gradients as that observed in our study’s high-intensity mowing plot. We encourage future research to test these predictions and continue to strengthen the evidence supporting the positive ecological effects of reduced lawn mowing.

### Limitations and future research

We have identified several avenues for refinement and future study through the course of this research. In addition to the site-level replication challenge outlined in the previous section, we draw attention to the potential of future studies to investigate the effects of reduced mowing over multiple years. Such datasets would enable implementation of multi-season dynamic occupancy models (Kéry and Schaub 2012), which have been used in related single-site, control-impact studies to assess how species richness and occupancy, but also demographic rates such as survival and colonisation, varied across years, yielding key insights into how quickly can insects in urban environments colonise new sites and whether they are likely to persist at the sites they colonise over multiple years (Mata et al. 2023). We also note that the short flowering period shown by the introduced forbs that came into flower during our study – likely influenced by lower-than-average rainfall – restricted our surveys of insect pollinators and other flower-visitor species to just two timepoints. While these datapoints still afforded us the opportunity to assess species richness and occupancy effects, it hindered our ability to generate properly replicated plant-pollinator interaction datasets and, therefore, to examine ideas around network structure and dynamics (Daniels et al. 2020, Silva et al. 2021, Mata et al. 2024). The prospect of collecting robust within-plot, multiple-site replicated plant-pollinator – as well as other bipartite and multi-trophic – interaction datasets for lawns experiencing reduced mowing serves as a continuous incentive for future research. Such studies would need to be carefully designed to meaningfully test community ecology hypotheses and demonstrate network typology effects (Dormann 2023).

Another core outcome of our study was achieving our overarching goal to work with the whole community of insects occurring in or visiting the experimental plots, which offered a more detailed understanding of the role that reduced mowing can play in providing food and habitat resources for assemblages of functionally similar species. A shortcoming of our study design, however, was the exclusion from our surveys of non-insect invertebrates. While we understood they would be theoretically present, they were intentionally omitted due to project scope constraints. Indeed, all surveyors anecdotally indicated observing many spider species falling in the sweep net during the low-intensity mowing plot surveys. Spiders comprise an abundant and diverse group of predatory species, including in urban environments (Willmott et al. 2025), and their absence in our dataset may help explain the marginally positive effect of vegetation height on the predatory assemblage. Yet another potential shortcoming, again due to project scope constraints, was our decision to not survey lepidopteran caterpillars, which would have enriched our findings for the herbivorous assemblage, and, if reared to maturity, allowed us to examine the effects of reduced mowing on the immature stages of moths and butterflies. Given the prevalence of parasitoid wasps in the low-intensity mowing plot, it is also likely that some of these caterpillars would have been parasitised, so examining parasitism and emergence rates would have provided an additional layer of understanding of the role that reduced mowing can play in supporting parasitoids and thus boosting the ecological functionality of lawns.

### Applications to lawn management

In a conscious attempt to bridge the gap between science and practice (Cadotte et al. 2017, Kurle et al. 2022), our study was specifically co-designed between local government practitioners and ecological researchers to synergistically generate scientific and practice insights that may be readily translated into actionable management innovations relevant both within and beyond the study context. As such, one of our study’s core desired outcomes was to establish pathways for collaborative teamwork between researchers, practitioners and other built-environment professionals to showcase how reduced lawn mowing can be a sound, evidence-based urban greening innovation that can be used across all levels of government and industry to meet local, regional and global biodiversity targets. As with many other urban greening and conservation innovations, however, uptake may be slow due to a series of risk, funding, knowledge, value alignment, and threat management implementation barriers (Soanes et al. 2023). We outline here examples of these barriers and propose potential solutions. While particular to our study context and sphere of local expertise, these barriers are likely to reflect similar circumstances present in urban environments globally, particularly given the ubiquity of lawns and the homogeneity of their management.

The first critical point to note revolves around management of noxious weeds – invasive, fast-growing plant species that outcompete indigenous species and that are targeted for mandatory control measures under national and state legislation (Commonwealth of Australia 2017) – especially those that emerge in response to reduced mowing. For instance, the widespread presence of the Chilean needle grass *Nasella neesiana* and serrated tussock *Nasella trichotoma* presents a distinct challenge for practitioners in the City of Merri-bek, with any future rollout of reduced mowing across the municipality likely required to be paired with strict, concurrent measures to control these weeds. While not officially declared as noxious, Kikuyu *Cenchrus clandestinus* – a rapidly spreading perennial grass indigenous to Africa, and often the dominant species in Australian lawns (Ignatieva et al. 2025) – was observed becoming overabundant towards the end of the austral autumn in a number of sites across the City of Merri-bek, suggesting that preventive management of any future low-mowing plots would be necessary to mitigate the establishment of Kikuyu monocultures, which we anticipate could severely dampen the positive effects of reduced mowing on insect communities. Regarding the potential critique that low-intensity mowing lawns may harbour problematic, invasive insect species, we highlight that, as other related studies within the same study area (Mata et al. 2021, 2023), we found a relatively low number of introduced species – representing only 3% and 14% of the total number of species observed in the sweep-netting and direct observation surveys, respectively – with none of them dominating in terms of their observed frequencies.

Another key challenge involves the interface between the low- and high-intensity mowing areas and its key role in communicating to greenspace users that the low-intensity mowing lawn is intentional rather than a result of neglect. The issue arises from the patchy, fluctuating nature of vegetation growth, which will vary at each site in response to a combination of factors, including soil legacy conditions (e.g. moisture, chemical and physical properties, seed bank) and seed dispersal potential. Uneven, inconsistent vegetation growth can mean that the desired clean and strong contrast between the low- and high-intensity mowing areas is not always attained, especially towards the end of mowing cycles. While failure to achieve a highly-contrasting interface – which should be achieved through a more frequent edge mowing regime targeted at retaining the boundary shape over time – would not affect the capacity of low-intensity mowing lawns to benefit insect communities and boost ecosystem functionality, it would certainly limit public acceptance of this management approach. Ensuring this outcome is essential, especially in light of other perceived risks (e.g. venomous snakes, fire load) that some greenspace users disproportionally associate with ‘long grass’. One potential solution to the interface challenge is to deliberately incorporate low-intensity mowing areas as design features of the greenspace, shaping the contrasting edges into aesthetically pleasing curves (Fig. S6). These areas could follow frequently used walking pathways and be complemented with educational signage to encourage insect observation. Finally, our co-designed approach has stimulated discussions around further ideas that could be examined in future research/practice partnerships. For instance, what is the optimal mowing frequency that maximises biodiversity gains while minimising weed dispersal? Would a mosaic design (i.e. maintaining lawn patches across the greenspace in various, differential growth stages) contribute to mitigate the detrimental effects likely to be experienced by the insect community when the low-intensity mowing areas are eventually mowed? From the outreach perspective, it would be interesting to examine how citizen science thinking and resources (Palma et al. 2024, Mason et al. 2025) may be operationalised to engage greenspace users and more broadly activate the community in understanding and advocating for the ecological benefits of reduced lawn mowing.

### Policy alignment and future outlook

Our study has advanced knowledge of the ecological benefits of reduced lawn mowing intensity and mapped a trajectory for stewards, practitioners and other built-environment professionals tasked with protecting existing nature or bringing nature back into their local urban areas, while concomitantly fulfilling broader mandates enshrined in regional and global policies. Indeed, the experimental component of our study was part of *The Lawn is Buzzing*, a project that was conducted at one of the demonstration sites within the network of low-intensity managed lawns established across the City of Merri-bek in 2023. This project was the direct result of a scoping review and community consultation campaign that provided background information and recommendations to Merri-bek City Council on actions with potential to promote insect biodiversity across the municipality (D. Echberg, unpublished data). As such, the study contributed to several of Merri-bek City Council’s policy goals, as outlined in the municipality’s Nature Plan (Merri-bek City Council 2020) and Open Space Strategy (Merri-bek City Council 2012), including: (1) addressing the problem of simplified landscapes, composed largely of trees and high-intensity mowed lawns, and their detrimental impacts on biodiversity; (2) making places for nature and increasing biodiversity through reframing lawns as potential habitat; (3) adopting up-to-date, sustainable environmental design in greenspace management; (4) engaging in collaborative work with research organisations to provide Council with the latest scientific evidence; (4) connecting people to nature through increasing understanding of the value of insects and the urban ecosystems that sustains them; and (5) seeking opportunities to make meaningful contributions to insect conservation.

In addition to advancing strategic policy goals, we see further value in this study – and any future broader rollout across Merri-bek and other municipalities – in its potential to challenge the dominant paradigm of what urban greenspaces should be. This includes inspiring people to see greenspaces as places for biodiversity as well as for people, and to question the long-standing aesthetic preference for ‘neat and tidy’ greenspaces. Cultivating ecological literacy widely and equitably across the community is a crucial step in this direction – from communicating fundamental ideas such as the value of biodiversity, the importance of insects for ecosystem functionality and the links between biodiversity and human health, through to perspective-shifting concepts such as untidy greenspaces being more ecologically rich than tidy ones. More distally, but fundamentally equally importantly, the study invites reflections on the broader sustainability and economic ramifications of reduced mowing; for example, the flow-on effects of reducing carbon emissions generated by mowing equipment and the freeing up of human and financial resources that could be allocated to more pressing ecological and social priorities.

## Acknowledgements

The authors would like to acknowledge the Traditional Custodians of the land and waterways on which this research took place and where we live and work, the Woi wurrung and Boon wurrung peoples of the eastern Kulin Nations – We pay our respects to their Elders, past, present and emerging, and honour their deep spiritual, cultural and customary connections to the land. This study was funded by Merri-bek City Council; we would like to extend our heartfelt thanks to Alistair Cowan, Tom Baldock and the Open Space Maintenance Crew Zone 8, including Crew Leader Danielle Violato and team members Sophie Merrett and Ben McKenzie, for their support and invaluable contributions to the study. Thanks to the iNaturalist community for providing species identifications, including *asiola*, Ben Parslow, *borisb*, Cecil Smith, Chris Lambkin, Dan Blamey, *gggpellas*, Jeong Yoo, Ken Harris, Kevin Williams, *lccg*, Matthew Connors, Nigel Main, oneanttofew, Otto Bell, Peter Ewin, Reiner Richter, Rod Lowther, Samuel Brown, Simon Taylor, Stephen Fricker, Susanna Heideman and Tony Daley. We would also like to acknowledge everyone that contributed to *The Little Things that Run the City* project, particularly to Amy Rogers, Lingna Zhang, Yvonne Lynch and Ian Shears who provided invaluable support to this project while working at The City of Melbourne.

## Authors’ contributions

All authors co-designed the project, with Luis Mata, Drew Echberg and Charlotte Napper contributing to the core project conceptualisation and all authors contributing to the project’s experimental design; Luis Mata, Drew Echberg and Charlotte Napper collected the data; Luis Mata and Estibaliz Palma organised and analysed the data, and conceptualised and developed visualisations and tables; all authors interpreted results; Drew Echberg and Charlotte Napper provided management recommendations and policy insights; Luis Mata led the writing of the manuscript; Drew Echberg, Charlotte Napper, Amy Hahs and Estibaliz Palma reviewed the manuscript and all authors approved it for publication.

## Conflict of Interest Statement

The authors declare that they have no known competing financial interests or personal relationships that could have appeared to influence the work reported in this paper.

## Open Research Statement

Data and codes to reproduce models and plots are already published and publicly available in Zenodo: https://doi.org/10.5281/zenodo.17181442.

## Notes

https://zenodo.org/records/17181442

### Competing Interest Statement

The authors have declared no competing interest.

