## Appendix S1 for "The Lawn is Buzzing: Increasing insect biodiversity in urban greenspaces through low-intensity mowing"

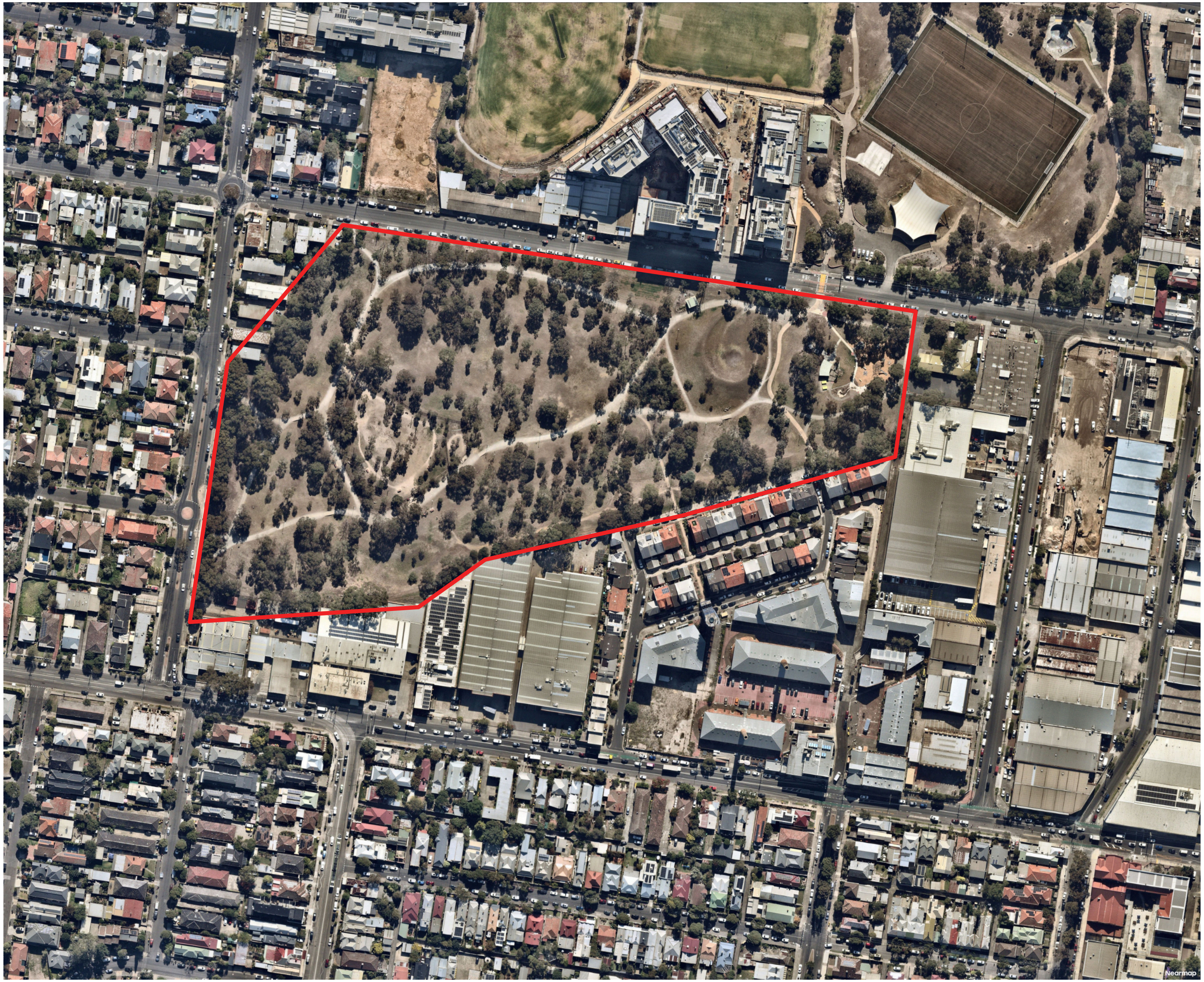

**Figure S1.** Aerial satellite image of Gilpin Park, Brunswick, Melbourne, Victoria, Australia and its surrounding suburban residential area. Photo provided by Nearmap as accessed through The University of Melbourne's library geospatial resources.

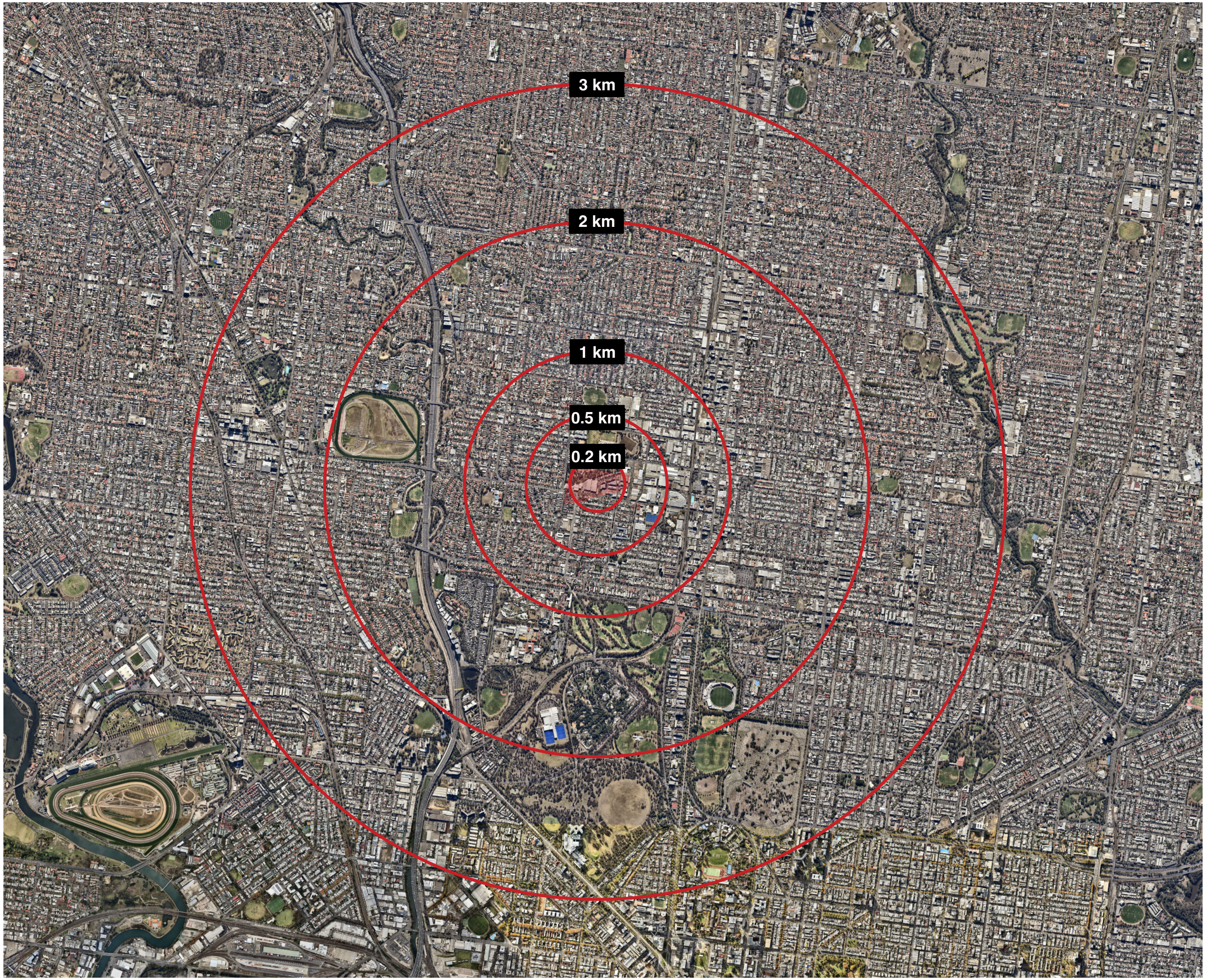


**Figure S2.** Aerial satellite image of the urban landscape surrounding Gilpin Park. Photo provided by Nearmap as accessed through The University of Melbourne's library geospatial resources.

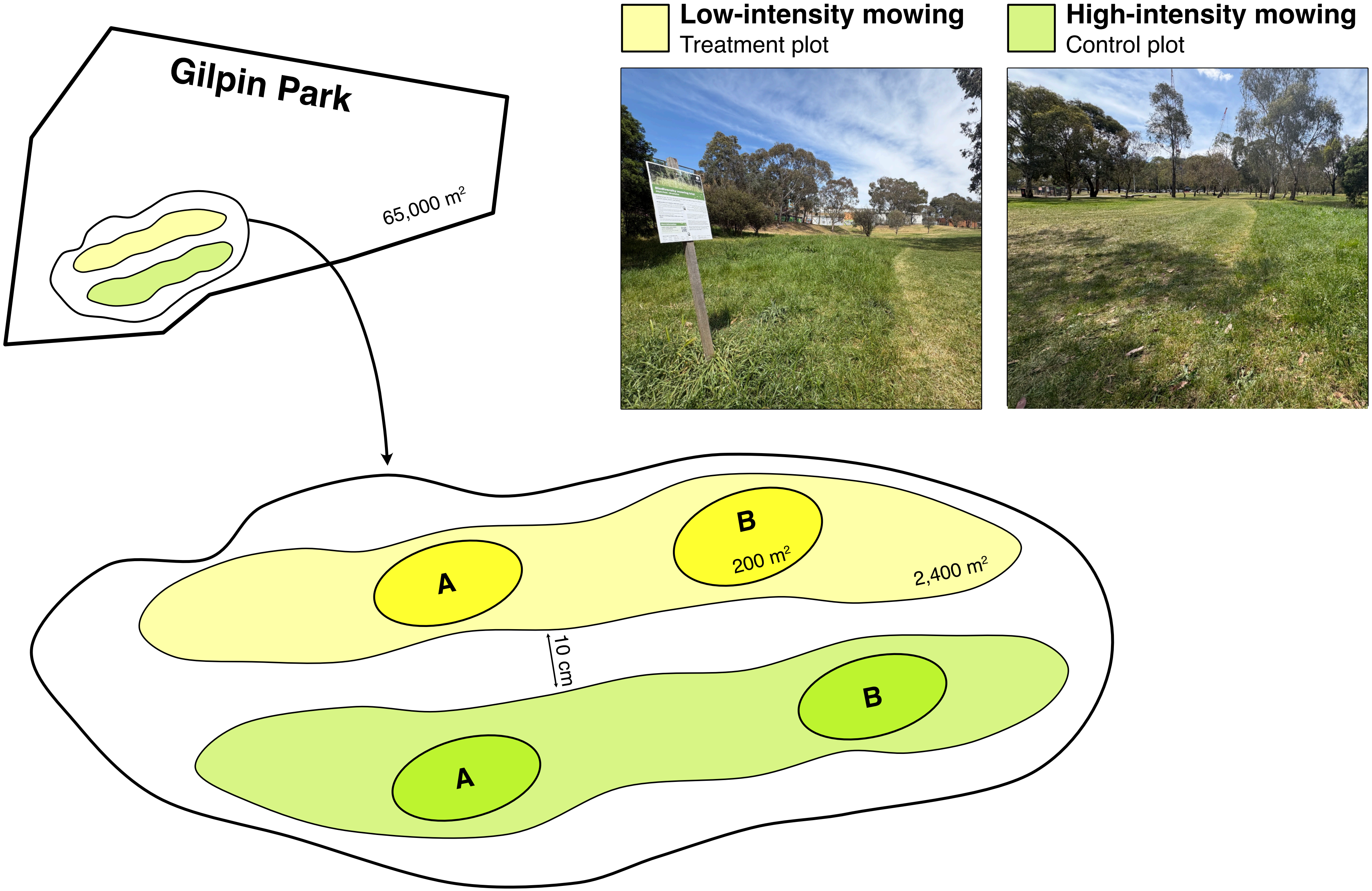


**Figure S3.** Experimental design of the study. Photos by Luis Mata.

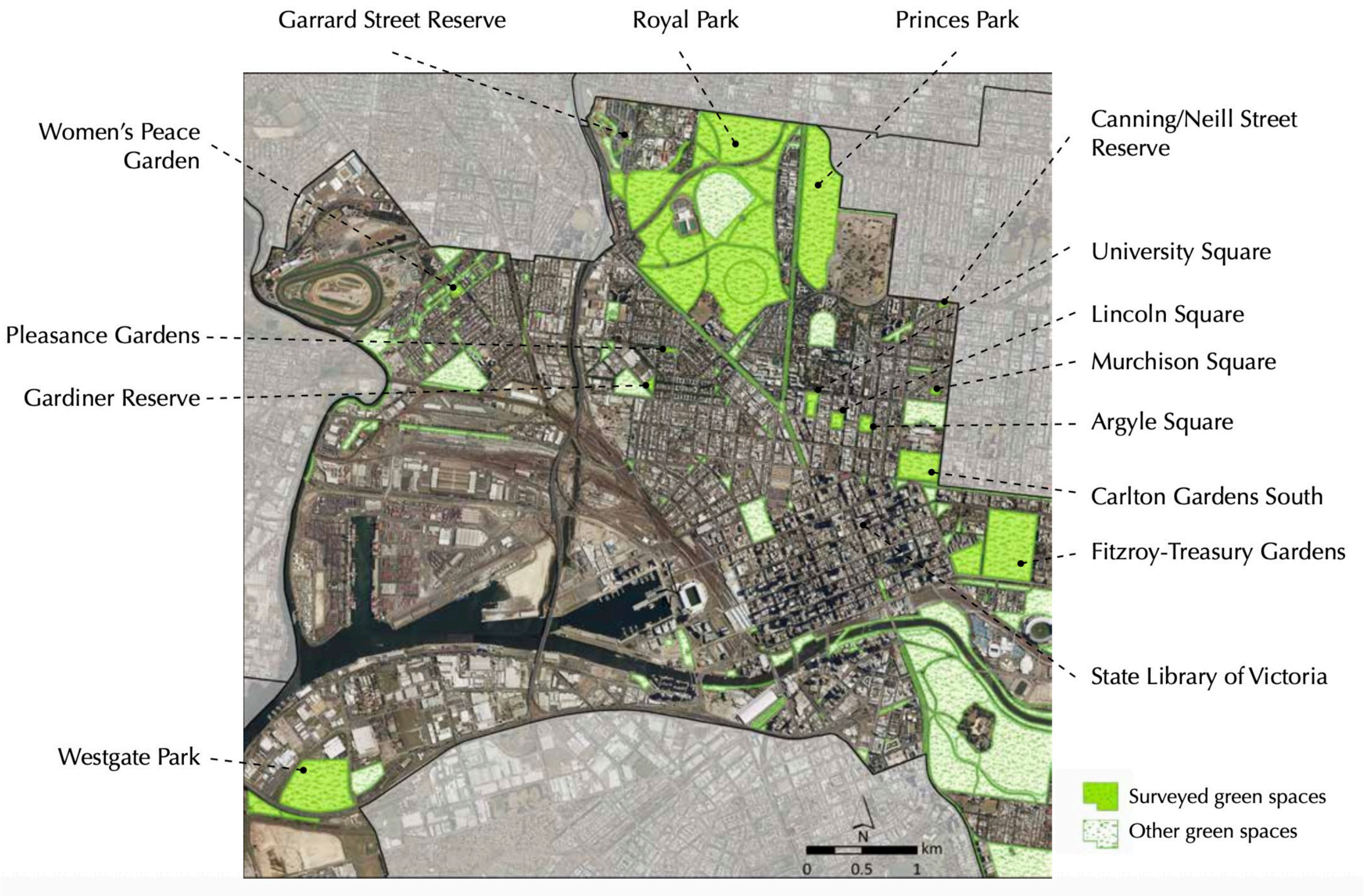


**Figure S4.** Location of the fifteen study sites surveyed within the City of Melbourne (Victoria, Australia) for *The Little Things that Run the City* project. Reproduced from Mata and colleagues (2016) *The Little Things that Run the City – Insect ecology, biodiversity and conservation in the City of Melbourne.*

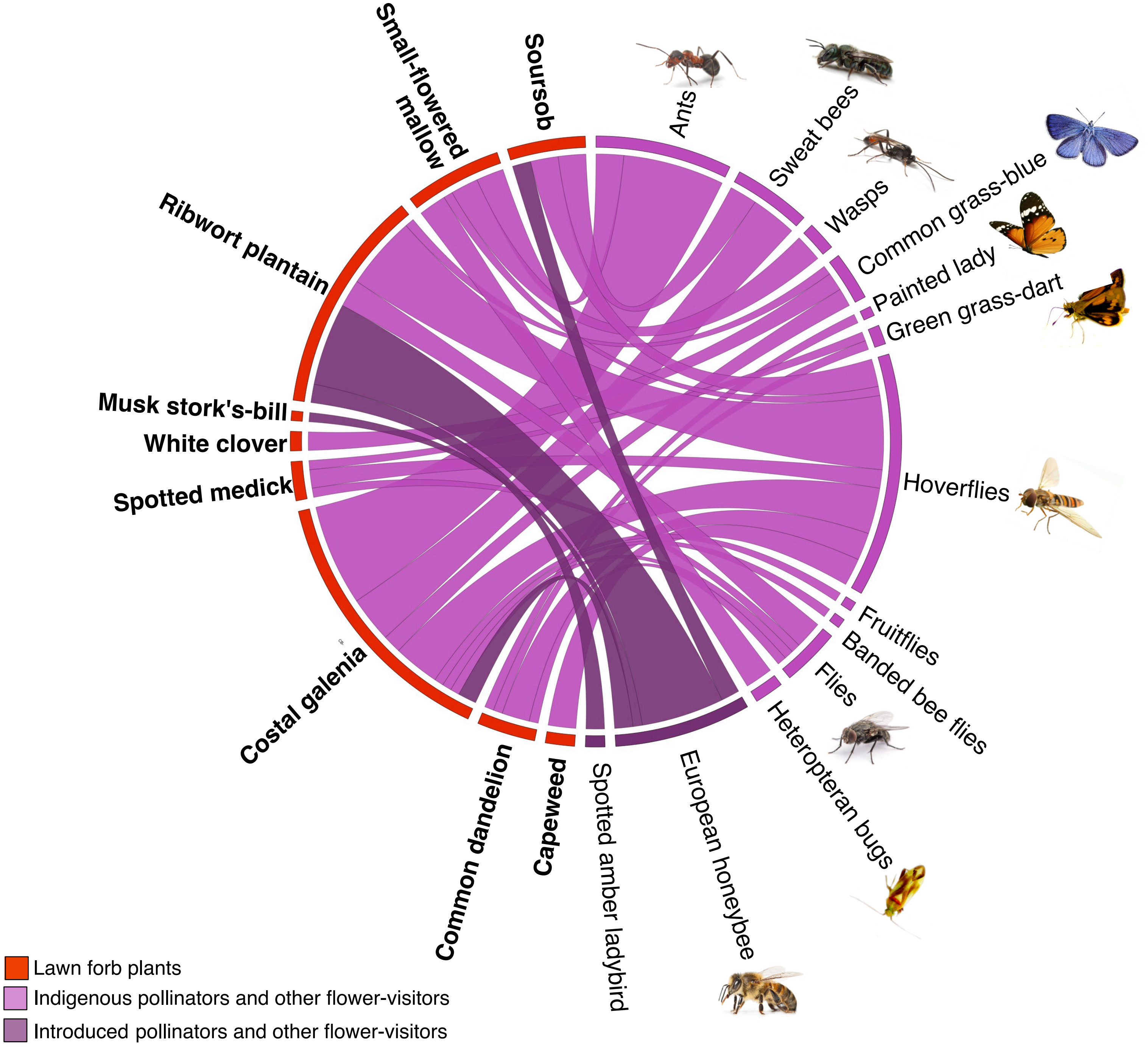


**Figure S5.** Quantitative network representation of the observed interaction between the flowering forb species (red boxes) and pollinator and other flower-visitor insect species (purple boxes) as recorded through the direct observations survey method. More details on the plant species are given in Table S2.

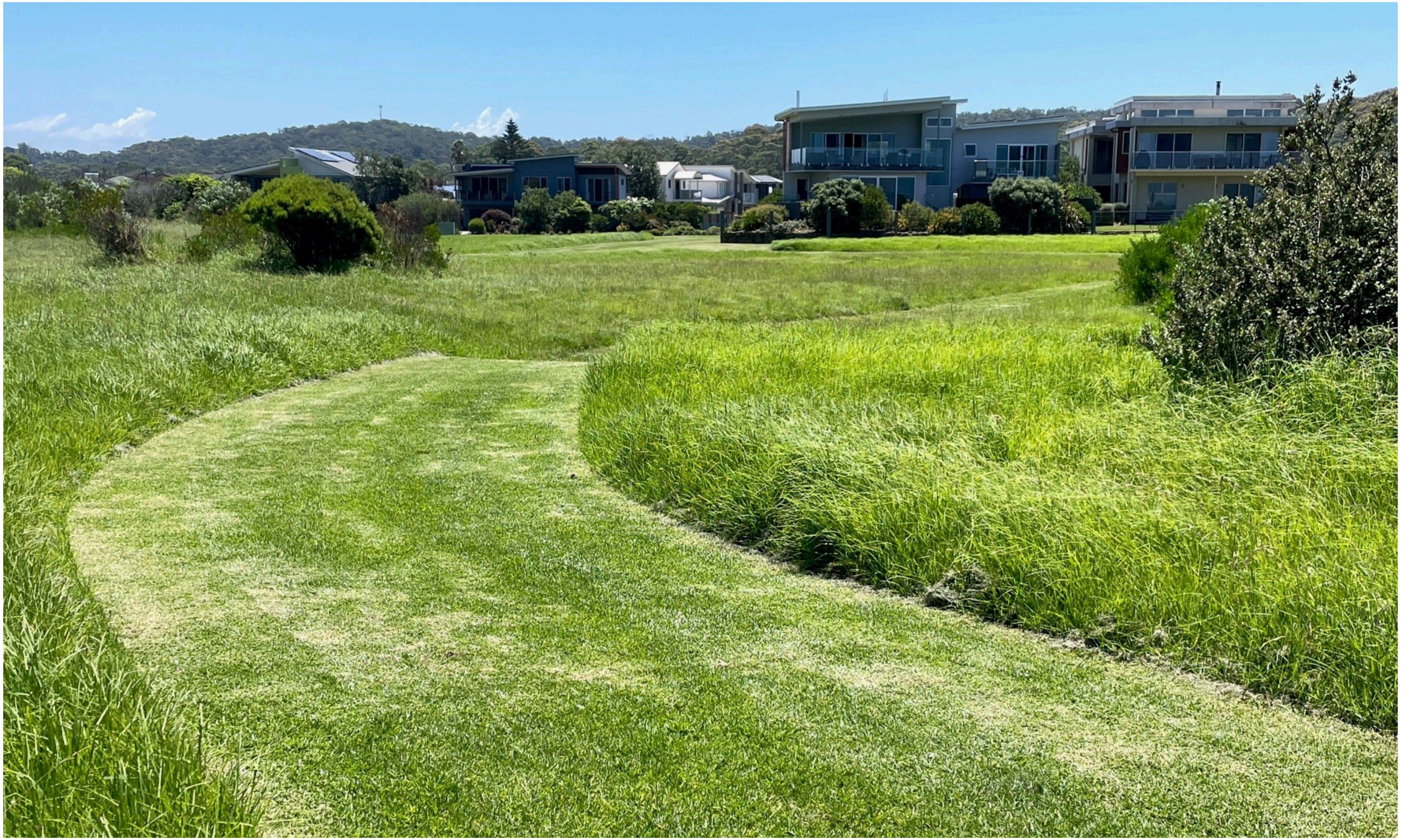

**Figure S6.** Example of how low-intensity mowing areas may be deliberately incorporated as a greenspace design feature by shaping the contrasting edges into aesthetically pleasing curves. Photo from Tomakin, New South Wales, Australia by Drew Echberg.

**Table S1.** The 194 insect species collected through sweep netting in *The Lawn is Buzzing* project in Gilpin Park, City of Merri-Bek (Melbourne, Victoria), their taxonomical associations, their functional group (DET: Detritivore; HER: Herbivore; PRE: Predator; PAR: Parasitoid), the treatment plot they were found at (High: high-intensity mowing; Low: low-intensity mowing), and the number of times they were recorded at each treatment. Introduced insect species are indicated with an *.

| Insect group I Species/morphospecies | Family | Functional group | | | | Mowing intensity | | High-intensity mowing plot | | Low-intensity mowing plot | |
|---|---|---|---|---|---|---|---|---|---|---|---|
| | | DET | HER | PRE | PAR | High | Low | Number of records (n=88) | % | Number of records (n=349) | % |
| **Ants [Hymenoptera: Apocrita: Aculeata: Formicoidea]** | | | | | | | | | | | |
| *Colobopsis sp. 1* | Formicidae | ■ | | | | ■ | | 1 | 1.1 | 0 | 0 |
| Formicidae 1 | Formicidae | ■ | | | | ■ | ■ | 7 | 8 | 10 | 2.9 |
| Formicidae 2 | Formicidae | ■ | | | | ■ | ■ | 1 | 1.1 | 1 | 0.3 |
| Formicidae 3 | Formicidae | ■ | | | | | ■ | 0 | 0 | 1 | 0.3 |
| Formicidae 4 | Formicidae | ■ | | | | | ■ | 0 | 0 | 1 | 0.3 |
| Formicidae 5 | Formicidae | ■ | | | | | ■ | 0 | 0 | 1 | 0.3 |
| *Pheidole sp. 1* | Formicidae | ■ | | | | ■ | ■ | 3 | 3.4 | 3 | 0.9 |
| Ponerinae 1 | Formicidae | ■ | | | | | ■ | 0 | 0 | 1 | 0.3 |
| **Bees [Hymenoptera: Apocrita: Aculeata: Apoidea: Anthophila]** | | | | | | | | | | | |
| *Apis mellifera* * | Apidae | | ■ | | | | ■ | 0 | 0 | 3 | 0.9 |
| *Lasioglossum sp. 1* | Halictidae | | ■ | | | | ■ | 0 | 0 | 1 | 0.3 |
| *Lasioglossum sp. 2* | Halictidae | | ■ | | | | ■ | 0 | 0 | 2 | 0.6 |
| *Lasioglossum sp. 3* | Halictidae | | ■ | | | ■ | ■ | 2 | 2.3 | 5 | 1.4 |
| *Lasioglossum sp. 4* | Halictidae | | ■ | | | | ■ | 0 | 0 | 1 | 0.3 |
| **Beetles [Coleoptera]** | | | | | | | | | | | |
| *Anthrenus sp. 1* * | Dermestidae | ■ | | | | ■ | | 1 | 1.1 | 0 | 0 |
| *Hippodamia variegata* * | Coccinellidae | | | ■ | | ■ | ■ | 1 | 1.1 | 2 | 0.6 |
| *Sitona discoideus* * | Curculionidae | | ■ | | | ■ | ■ | 2 | 2.3 | 8 | 2.3 |
| Alticini 1 | Chrysomelidae | | ■ | | | ■ | ■ | 1 | 1.1 | 3 | 0.9 |
| Anthicidae 1 | Anthicidae | ■ | | | | | ■ | 0 | 0 | 1 | 0.3 |
| Anthicidae 2 | Anthicidae | ■ | | | | | ■ | 0 | 0 | 1 | 0.3 |
| Anthicidae 3 | Anthicidae | ■ | | | | | ■ | 0 | 0 | 2 | 0.6 |
| Anthicidae 4 | Anthicidae | ■ | | | | | ■ | 0 | 0 | 1 | 0.3 |
| Anthicidae 5 | Anthicidae | ■ | | | | | ■ | 0 | 0 | 1 | 0.3 |
| Apionini 1 | Brentidae | | ■ | | | | ■ | 0 | 0 | 1 | 0.3 |
| Baridinae 1 | Curculionidae | | ■ | | | | ■ | 0 | 0 | 2 | 0.6 |
| Carabidae 1 | Carabidae | | | ■ | | ■ | | 1 | 1.1 | 0 | 0 |
| Carabidae 2 | Carabidae | | | ■ | | | ■ | 0 | 0 | 1 | 0.3 |
| *Chauliognathus lugubris* | Cantharidea | | | ■ | | | ■ | 0 | 0 | 1 | 0.3 |
| Chrysomelidae 1 | Chrysomelidae | | ■ | | | | ■ | 0 | 0 | 1 | 0.3 |
| *Coccinella transversalis* | Coccinellidae | | | ■ | | ■ | ■ | 1 | 1.1 | 2 | 0.6 |
| Coccinellidae 1 | Coccinellidae | | | ■ | | | ■ | 0 | 0 | 1 | 0.3 |
| Coccinellidae 2 | Coccinellidae | | | ■ | | | ■ | 0 | 0 | 1 | 0.3 |
| *Cortinicara sp.1* | Latriidae | ■ | | | | ■ | ■ | 4 | 4.5 | 10 | 2.9 |

**Table S1.** ***Cont.***

| Insect group I Species/morphospecies | Family | Functional group | | | | Mowing intensity | | High-intensity mowing plot | | Low-intensity mowing plot | |
|---|---|---|---|---|---|---|---|---|---|---|---|
| | | DET | HER | PRE | PAR | High | Low | Number of records (n=88) | % | Number of records (n=349) | % |
| **Beetles *Cont.*** | | | | | | | | | | | |
| *Cryptocephalini 1* | Chrysomelidae | | ■ | | | | ■ | 0 | 0 | 1 | 0.3 |
| *Diomus notescens* | Coccinellidae | | | ■ | | | ■ | 0 | 0 | 2 | 0.6 |
| *Diomus sp. 1* | Coccinellidae | | | ■ | | | ■ | 0 | 0 | 1 | 0.3 |
| Elateridae 1 | Elateridae | ■ | | | | | ■ | 0 | 0 | 1 | 0.3 |
| *Euciodes suturalis* | Anthribidae | | ■ | | | | ■ | 0 | 0 | 1 | 0.3 |
| Mordellidae 1 | Mordellidae | ■ | | | | | ■ | 0 | 0 | 1 | 0.3 |
| Mordellidae 2 | Mordellidae | ■ | | | | | ■ | 0 | 0 | 1 | 0.3 |
| Opatrini 1 | Tenebrionidae | ■ | | | | ■ | | 1 | 1.1 | 0 | 0 |
| *Paederus sp. 1* | Staphylinidae | | | ■ | | | ■ | 0 | 0 | 1 | 0.3 |
| Phalacridae 1 | Phalacridae | | ■ | | | | ■ | 0 | 0 | 1 | 0.3 |
| Polyphaga 1 | | | ■ | | | | ■ | 0 | 0 | 1 | 0.3 |
| Polyphaga 2 | | | ■ | | | ■ | | 1 | 1.1 | 0 | 0 |
| *Rhinoncus australis* | Curculionidae | | ■ | | | | ■ | 0 | 0 | 1 | 0.3 |
| **Booklice [Psocoptera]** | | | | | | | | | | | |
| Psocoptera 1 | | ■ | | | | | ■ | 0 | 0 | 1 | 0.3 |
| **Brachyceran flies [Diptera: Brachycera]** | | | | | | | | | | | |
| Anabarhynchus sp. 1 | Therevidae | | | ■ | | | ■ | 0 | 0 | 1 | 0.3 |
| Anthomyia punctipennis | Anthomyiidae | ■ | | | | ■ | ■ | 2 | 2.3 | 2 | 0.6 |
| Anthomyiidae 1 | Anthomyiidae | ■ | | | | ■ | ■ | 1 | 1.1 | 1 | 0.3 |
| Anthomyiidae 2 | Anthomyiidae | ■ | | | | ■ | ■ | 3 | 3.4 | 6 | 1.7 |
| *Atherigona sp. 1* | Muscidae | ■ | | | | | ■ | 0 | 0 | 2 | 0.6 |
| *Australoactina sp. 1* | Stratiomyidae | ■ | | | | | ■ | 0 | 0 | 2 | 0.6 |
| *Austrotephritis poenia* | Tephritidae | | ■ | | | ■ | ■ | 1 | 1.1 | 2 | 0.6 |
| Brachycera 1 | | ■ | | | | ■ | ■ | 4 | 4.5 | 5 | 1.4 |
| Brachycera 2 | | ■ | | | | | ■ | 0 | 0 | 1 | 0.3 |
| Brachycera 3 | | ■ | | | | ■ | ■ | 1 | 1.1 | 3 | 0.9 |
| Brachycera 4 | | ■ | | | | ■ | ■ | 1 | 1.1 | 1 | 0.3 |
| Brachycera 5 | | ■ | | | | ■ | ■ | 1 | 1.1 | 1 | 0.3 |
| Brachycera 6 | | ■ | | | | ■ | | 1 | 1.1 | 0 | 0 |
| Brachycera 7 | | ■ | | | | ■ | ■ | 2 | 2.3 | 2 | 0.6 |
| Brachycera 8 | | ■ | | | | | ■ | 0 | 0 | 2 | 0.6 |
| Brachycera 9 | | ■ | | | | | ■ | 0 | 0 | 1 | 0.3 |
| Brachycera 10 | | ■ | | | | | ■ | 0 | 0 | 4 | 1.1 |
| Brachycera 11 | | ■ | | | | | ■ | 0 | 0 | 1 | 0.3 |
| Brachycera 12 | | ■ | | | | ■ | | 1 | 1.1 | 0 | 0 |
| Brachycera 13 | | ■ | | | | | ■ | 0 | 0 | 1 | 0.3 |

**Table S1.** ***Cont.***

| Insect group I Species/morphospecies | Family | Functional group | | | | Mowing intensity | | High-intensity mowing plot | | Low-intensity mowing plot | |
|---|---|---|---|---|---|---|---|---|---|---|---|
| | | DET | HER | PRE | PAR | High | Low | Number of records (n=88) | % | Number of records (n=349) | % |
| **Brachyceran flies *Cont.*** | | | | | | | | | | | |
| *Cerdistus sp. 1* | Asilidae | | | ■ | | | ■ | 0 | 0 | 1 | 0.3 |
| *Coenosia sp. 1* | Muscidae | | | ■ | | ■ | ■ | 4 | 4.5 | 10 | 2.9 |
| *Depressa atrata* | Lauxaniidae | ■ | | | | | ■ | 0 | 0 | 1 | 0.3 |
| Diaphorinae 1 | Dolichopodidae | | | ■ | | | ■ | 0 | 0 | 1 | 0.3 |
| *Dichetophora sp. 1* | Sciomyzidae | | | ■ | | | ■ | 0 | 0 | 3 | 0.9 |
| Dolichopodidae 1 | Dolichopodidae | | | ■ | | | ■ | 0 | 0 | 1 | 0.3 |
| Hybotidae 1 | Hybotidae | ■ | | | | | ■ | 0 | 0 | 1 | 0.3 |
| *Hydrellia tritici* | Ephydridae | | ■ | | | ■ | ■ | 9 | 10.2 | 10 | 2.9 |
| Muscidae 1 | Muscidae | ■ | | | | | ■ | 0 | 0 | 3 | 0.9 |
| Muscidae 2 | Muscidae | ■ | | | | ■ | ■ | 1 | 1.1 | 2 | 0.6 |
| Oestroidea 1 | | ■ | | | | ■ | ■ | 1 | 1.1 | 1 | 0.3 |
| Oestroidea 2 | | ■ | | | | | ■ | 0 | 0 | 1 | 0.3 |
| Oestroidea 3 | | ■ | | | | ■ | | 1 | 1.1 | 0 | 0 |
| Orthorrhapha 1 | | ■ | | | | ■ | | 1 | 1.1 | 0 | 0 |
| *Parapalaeosepsis plebeia* | Sepsidae | ■ | | | | ■ | ■ | 2 | 2.3 | 10 | 2.9 |
| *Phasia sp. 1* | Tachinidae | | | | ■ | | ■ | 0 | 0 | 1 | 0.3 |
| *Poecilohetaerus aquilus* | Lauxaniidae | ■ | | | | ■ | ■ | 3 | 3.4 | 8 | 2.3 |
| *Rivellia sp. 1* | Platystomatidae | ■ | | | | | ■ | 0 | 0 | 1 | 0.3 |
| *Sapromyza mallochiana* | Lauxaniidae | ■ | | | | | ■ | 0 | 0 | 4 | 1.1 |
| *Simosyrphus grandicornis* | Syrphidae | | | ■ | | ■ | ■ | 1 | 1.1 | 1 | 0.3 |
| *Spathulina acroleuca* | Tephritidae | | ■ | | | ■ | ■ | 1 | 1.1 | 1 | 0.3 |
| *Sphenella ruficeps* | Tephritidae | | ■ | | | | ■ | 0 | 0 | 2 | 0.6 |
| *Steganopsis melanogaster* | Lauxaniidae | ■ | | | | ■ | ■ | 1 | 1.1 | 6 | 1.7 |
| Therevidae 1 | Therevidae | | | ■ | | | ■ | 0 | 0 | 1 | 0.3 |
| **Earwigs [Dermaptera]** | | | | | | | | | | | |
| *Forficula dentata* * | Forficulidae | | ■ | | | | ■ | 0 | 0 | 3 | 0.9 |
| **Grasshoppers, crickets and katydids [Orthoptera]** | | | | | | | | | | | |
| Catantopini 1 | Acrididae | | ■ | | | | ■ | 0 | 0 | 1 | 0.3 |
| *Conocephalus albescens* | Tettigoniidae | | ■ | | | | ■ | 0 | 0 | 1 | 0.3 |
| *Conocephalus upoluensis* | Tettigoniidae | | ■ | | | | ■ | 0 | 0 | 1 | 0.3 |
| Gryllidae 1 | Gryllidae | | ■ | | | | ■ | 0 | 0 | 1 | 0.3 |
| *Schizobothrus flavovittatus* | Acrididae | | ■ | | | | ■ | 0 | 0 | 3 | 0.9 |
| Trigonidiinae 1 | Trigonidiidae | | ■ | | | | ■ | 0 | 0 | 2 | 0.6 |

**Table S1.** ***Cont.***

| Insect group I Species/morphospecies | Family | Functional group: DET | HER | PRE | PAR | Mowing intensity: High | Low | High-intensity mowing plot: Number of records (n=88) | % | Low-intensity mowing plot: Number of records (n=349) | % |
|---|---|---|---|---|---|---|---|---|---|---|---|
| **Heteropteran bugs [Hemiptera: Heteroptera]** | | | | | | | | | | | |
| *Pyrrhocoris apterus* * | Pyrrocoridae | | ■ | | | | ■ | 0 | 0 | 3 | 0.9 |
| *Amorbus sp. 1* | Coreidae | | ■ | | | | ■ | 0 | 0 | 1 | 0.3 |
| *Cermatulus nasalis* | Pentatomidae | | ■ | | | | ■ | 0 | 0 | 1 | 0.3 |
| Cydnidae 1 | Cydnidae | | ■ | | | | ■ | 0 | 0 | 1 | 0.3 |
| *Dindymus versicolor* | Pyrrocoridae | | ■ | | | | ■ | 0 | 0 | 2 | 0.6 |
| *Eribotes sp. 2* | Pentatomidae | | ■ | | | | ■ | 0 | 0 | 1 | 0.3 |
| *Melanacanthus sp. 1* | Alydidae | | ■ | | | | ■ | 0 | 0 | 1 | 0.3 |
| Miridae 1 | Miridae | | ■ | | | | ■ | 0 | 0 | 1 | 0.3 |
| Miridae 2 | Miridae | | ■ | | | | ■ | 0 | 0 | 1 | 0.3 |
| *Mutsca brevicornis* | Alydidae | | ■ | | | | ■ | 0 | 0 | 1 | 0.3 |
| *Nabis kinbergii* | Nabidae | | | ■ | | | ■ | 0 | 0 | 5 | 1.4 |
| *Nysius vinitor* | Lygaeidae | | ■ | | | ■ | ■ | 1 | 1.1 | 3 | 0.9 |
| *Oxycarenus luctuosus* | Oxycarenidae | | ■ | | | | ■ | 0 | 0 | 1 | 0.3 |
| Pentatomidae 1 | Pentatomidae | | ■ | | | | ■ | 0 | 0 | 1 | 0.3 |
| *Sidnia kinbergi* | Miridae | | ■ | | | | ■ | 0 | 0 | 1 | 0.3 |
| *Stenophylla macreta* | Pachygronthidae | | ■ | | | | ■ | 0 | 0 | 3 | 0.9 |
| Thaumastocoridae 1 | Thaumastocoridae | | ■ | | | | ■ | 0 | 0 | 1 | 0.3 |
| **Jumping plant lice [Hemiptera: Sternorrhyncha: Psylloidea]** | | | | | | | | | | | |
| Psylloidea 1 | | | ■ | | | | ■ | 0 | 0 | 1 | 0.3 |
| Psylloidea 2 | | | ■ | | | | ■ | 0 | 0 | 1 | 0.3 |
| Psylloidea 3 | | | ■ | | | ■ | | 1 | 1.1 | 0 | 0 |
| **Lacewings [Neuroptera]** | | | | | | | | | | | |
| *Micromus tasmaniae* | Hemerobiidae | | | ■ | | | ■ | 0 | 0 | 5 | 1.4 |
| **Leafhoppers [Hemiptera: Auchenorrhyncha: Membracoidea]** | | | | | | | | | | | |
| *Brunotartessus fulvus* | Cicadellidae | | ■ | | | ■ | | 1 | 1.1 | 0 | 0 |
| Cicadellidae 1 | Cicadellidae | | ■ | | | ■ | ■ | 3 | 3.4 | 2 | 0.6 |
| Cicadellidae 2 | Cicadellidae | | ■ | | | | ■ | 0 | 0 | 2 | 0.6 |
| Cicadellidae 3 | Cicadellidae | | ■ | | | ■ | | 1 | 1.1 | 0 | 0 |
| Cicadellidae 4 | Cicadellidae | | ■ | | | ■ | ■ | 1 | 1.1 | 2 | 0.6 |
| Cicadellidae 5 | Cicadellidae | | ■ | | | ■ | ■ | 1 | 1.1 | 4 | 1.1 |
| Cicadellidae 6 | Cicadellidae | | ■ | | | | ■ | 0 | 0 | 1 | 0.3 |
| Cicadellidae 7 | Cicadellidae | | ■ | | | ■ | ■ | 1 | 1.1 | 3 | 0.9 |
| Cicadellidae 8 | Cicadellidae | | ■ | | | | ■ | 0 | 0 | 2 | 0.6 |
| Cicadellidae 9 | Cicadellidae | | ■ | | | ■ | ■ | 1 | 1.1 | 2 | 0.6 |

Table S1. *Cont.*

| Insect group I Species/morphospecies | Family | Functional group | | | | Mowing intensity | | High-intensity mowing plot | | Low-intensity mowing plot | |
|---|---|---|---|---|---|---|---|---|---|---|---|
| | | DET | HER | PRE | PAR | High | Low | Number of records (n=88) | % | Number of records (n=349) | % |
| **Leafhoppers *Cont.*** | | | | | | | | | | | |
| Cicadellidae 10 | Cicadellidae | | ■ | | | | ■ | 0 | 0 | 1 | 0.3 |
| Cicadellidae 11 | Cicadellidae | | ■ | | | | ■ | 0 | 0 | 1 | 0.3 |
| Cicadellidae 12 | Cicadellidae | | ■ | | | | ■ | 0 | 0 | 1 | 0.3 |
| Cicadellidae 13 | Cicadellidae | | ■ | | | | ■ | 0 | 0 | 1 | 0.3 |
| Cicadellidae 14 | Cicadellidae | | ■ | | | | ■ | 0 | 0 | 1 | 0.3 |
| Cicadellidae 15 | Cicadellidae | | ■ | | | | ■ | 0 | 0 | 1 | 0.3 |
| **Nematoceran flies [Diptera: Nematocera]** | | | | | | | | | | | |
| Tipulomorpha 1 | | ■ | | | | | ■ | 0 | 0 | 1 | 0.3 |
| **Parasitoid wasps [Hymenoptera: Apocrita: Parasitica]** | | | | | | | | | | | |
| Alysiinae 1 | Braconidae | | | | ■ | | ■ | 0 | 0 | 1 | 0.3 |
| Braconidae 1 | Braconidae | | | | ■ | | ■ | 0 | 0 | 2 | 0.6 |
| Braconidae 2 | Braconidae | | | | ■ | ■ | ■ | 1 | 1.1 | 2 | 0.6 |
| Braconidae 3 | Braconidae | | | | ■ | | ■ | 0 | 0 | 1 | 0.3 |
| Braconidae 4 | Braconidae | | | | ■ | | ■ | 0 | 0 | 2 | 0.6 |
| Braconidae 5 | Braconidae | | | | ■ | ■ | ■ | 1 | 1.1 | 1 | 0.3 |
| Campopleginae 1 | Ichneumonidae | | | | ■ | | ■ | 0 | 0 | 2 | 0.6 |
| Chalcididae 1 | Chalcididae | | | | ■ | | ■ | 0 | 0 | 1 | 0.3 |
| Chalcidoidea 1 | | | | | ■ | | ■ | 0 | 0 | 1 | 0.3 |
| Chalcidoidea 2 | | | | | ■ | | ■ | 0 | 0 | 1 | 0.3 |
| Chalcidoidea 3 | | | | | ■ | | ■ | 0 | 0 | 2 | 0.6 |
| Cheloninae 1 | Braconidae | | | | ■ | | ■ | 0 | 0 | 2 | 0.6 |
| Cheloninae 2 | Braconidae | | | | ■ | ■ | ■ | 1 | 1.1 | 1 | 0.3 |
| Cheloninae 3 | Braconidae | | | | ■ | | ■ | 0 | 0 | 1 | 0.3 |
| *Enicospilus sp. 1* | Ichneumonidae | | | | ■ | | ■ | 0 | 0 | 1 | 0.3 |
| *Epitranus sp. 1* | Chalcididae | | | | ■ | | ■ | 0 | 0 | 1 | 0.3 |
| *Gasteruption raphidioides* | Gasteruptiidae | | | | ■ | | ■ | 0 | 0 | 1 | 0.3 |
| Gravenhorstiini 1 | Ichneumonidae | | | | ■ | | ■ | 0 | 0 | 1 | 0.3 |
| Ichneumonoidea 1 | | | | | ■ | | ■ | 0 | 0 | 3 | 0.9 |
| *Lissonota sp. 1* | Ichneumonidae | | | | ■ | | ■ | 0 | 0 | 2 | 0.6 |
| *Myrmeleonostenus sp. 1* | Ichneumonidae | | | | ■ | | ■ | 0 | 0 | 2 | 0.6 |

**Table S1.** ***Cont.***

| Insect group I Species/morphospecies | Family | Functional group: DET | HER | PRE | PAR | Mowing intensity: High | Low | High-intensity mowing plot: Number of records (n=88) | % | Low-intensity mowing plot: Number of records (n=349) | % |
|---|---|---|---|---|---|---|---|---|---|---|---|
| **Parasitoid wasps** ***Cont.*** | | | | | | | | | | | |
| Parasitica 1 | | | | | ■ | ■ | | 1 | 1.1 | 0 | 0 |
| Parasitica 2 | | | | | ■ | | ■ | 0 | 0 | 1 | 0.3 |
| Parasitica 3 | | | | | ■ | | ■ | 0 | 0 | 3 | 0.9 |
| Parasitica 4 | | | | | ■ | | ■ | 0 | 0 | 3 | 0.9 |
| Parasitica 5 | | | | | ■ | | ■ | 0 | 0 | 1 | 0.3 |
| Parasitica 6 | | | | | ■ | | ■ | 0 | 0 | 2 | 0.6 |
| Parasitica 7 | | | | | ■ | | ■ | 0 | 0 | 1 | 0.3 |
| Parasitica 8 | | | | | ■ | | ■ | 0 | 0 | 1 | 0.3 |
| Parasitica 9 | | | | | ■ | | ■ | 0 | 0 | 1 | 0.3 |
| Parasitica 10 | | | | | ■ | ■ | ■ | 1 | 1.1 | 3 | 0.9 |
| Parasitica 11 | | | | | ■ | | ■ | 0 | 0 | 1 | 0.3 |
| Parasitica 12 | | | | | ■ | | ■ | 0 | 0 | 3 | 0.9 |
| Parasitica 13 | | | | | ■ | ■ | ■ | 1 | 1.1 | 2 | 0.6 |
| Parasitica 14 | | | | | ■ | | ■ | 0 | 0 | 2 | 0.6 |
| Parasitica 15 | | | | | ■ | | ■ | 0 | 0 | 1 | 0.3 |
| Parasitica 16 | | | | | ■ | | ■ | 0 | 0 | 2 | 0.6 |
| Parasitica 17 | | | | | ■ | | ■ | 0 | 0 | 1 | 0.3 |
| Parasitica 18 | | | | | ■ | | ■ | 0 | 0 | 2 | 0.6 |
| Parasitica 19 | | | | | ■ | | ■ | 0 | 0 | 1 | 0.3 |
| Parasitica 20 | | | | | ■ | | ■ | 0 | 0 | 1 | 0.3 |
| Parasitica 21 | | | | | ■ | | ■ | 0 | 0 | 1 | 0.3 |
| Parasitica 22 | | | | | ■ | | ■ | 0 | 0 | 1 | 0.3 |
| Parasitica 23 | | | | | ■ | | ■ | 0 | 0 | 1 | 0.3 |
| Parasitica 24 | | | | | ■ | | ■ | 0 | 0 | 1 | 0.3 |
| Parasitica 25 | | | | | ■ | | ■ | 0 | 0 | 1 | 0.3 |
| *Setanta sp. 1* | Ichneumonidae | | | | ■ | | ■ | 0 | 0 | 1 | 0.3 |
| Tersilochinae 1 | Ichneumonidae | | | | ■ | | ■ | 0 | 0 | 1 | 0.3 |
| **Planthoppers [Hemiptera: Auchenorryncha: Fulgoromorpha]** | | | | | | | | | | | |
| *Anzora unicolor* | Flatidae | | ■ | | | | ■ | 0 | 0 | 1 | 0.3 |
| Delphacidae 1 | Delphacidae | | ■ | | | | ■ | 0 | 0 | 4 | 1.1 |
| Delphacidae 2 | Delphacidae | | ■ | | | | ■ | 0 | 0 | 1 | 0.3 |
| Delphacidae 3 | Delphacidae | | ■ | | | | ■ | 0 | 0 | 1 | 0.3 |

**Table S1.** ***Cont.***

| Insect group I Species/morphospecies | Family | Functional group | | | | Mowing intensity | | High-intensity mowing plot | | Low-intensity mowing plot | |
|---|---|---|---|---|---|---|---|---|---|---|---|
| | | DET | HER | PRE | PAR | High | Low | Number of records (n=88) | % | Number of records (n=349) | % |
| **Stinging wasps [Hymenoptera: Apocrita: Aculeata, excl. Apoidea and Formicoidea]** | | | | | | | | | | | |
| Bethylidae 1 | Bethylidae | | | | ■ | | ■ | 0 | 0 | 1 | 0.3 |
| Bethylidae 2 | Bethylidae | | | | ■ | | ■ | 0 | 0 | 1 | 0.3 |
| Bethylidae 3 | Bethylidae | | | | ■ | ■ | ■ | 1 | 1.1 | 4 | 1.1 |
| Dasymutillini 1 | Mutillidae | | ■ | | | | ■ | 0 | 0 | 1 | 0.3 |
| Pompilidae 1 | Pompilidae | | | | ■ | | ■ | 0 | 0 | 1 | 0.3 |
| Pompilidae 2 | Pompilidae | | | | ■ | | ■ | 0 | 0 | 1 | 0.3 |
| Thynnidae 1 | Thynnidae | | | | ■ | | ■ | 0 | 0 | 1 | 0.3 |

**Table S2.** Plant species surveyed through direct observations in *The Lawn is Buzzing project* in Gilpin Park, Merri-Bek (Melbourne, Victoria), their taxonomical associations, native range, the treatment plot they were found at (High: high-intensity mowing; Low: low-intensity mowing), the number of surveys conducted on them for each treatment, and their overall estimated insect species richness. All species are introduced to Australia.

| Common name | Scientific name | Family | Order | Native range | Mowing intensity | | High-intensity mowing plot | | Low-intensity mowing plot | | Species richness | | | |
|---|---|---|---|---|---|---|---|---|---|---|---|---|---|---|
| | | | | | High | Low | Number of surveys (n=5) | % | Number of surveys (n=19) | % | Mean | SD | 2.5% | 97.5% |
| Coastal galenia | *Aizoon pubescens* | Aizoaceae | Caryophyllales | South Africa | ■ | ■ | 1 | 20.0 | 5 | 26.3 | 4.2 | 0.2 | 4.0 | 4.7 |
| Capeweed | *Arctotheca calendula* | Asteraceae | Asterales | South Africa | ■ | | 1 | 20.0 | 0 | 0 | 0.0 | 0.0 | 0.0 | 0.0 |
| Musk strok's-bill | *Erodium moschatum* | Geraniaceae | Geraniales | Asia, Europe, Northern Africa | | ■ | 0 | 0.0 | 1 | 5.3 | 1.1 | 0.3 | 1.0 | 2.0 |
| Small-flowered mallow | *Malva parviflora* | Malvaceae | Malvales | Asia, Europe, Northern Africa | ■ | ■ | 1 | 20.0 | 2 | 10.5 | 2.2 | 0.3 | 2.0 | 3.0 |
| Spotted medick | *Medicago arabica* | Fabaceae | Fabales | Asia, Europe, Northern Africa | | ■ | 0 | 0.0 | 2 | 10.5 | 1.3 | 0.4 | 1.0 | 2.0 |
| Soursob | *Oxalis pes-caprae* | Oxalidaceae | Oxalidales | South Africa | | ■ | 0 | 0.0 | 1 | 5.3 | 3.3 | 0.4 | 3.0 | 4.0 |
| Ribwort plantain | *Plantago lanceolata* | Plantaginaceae | Lamiales | Europe | ■ | ■ | 1 | 20.0 | 7 | 36.8 | 1.5 | 0.2 | 1.3 | 2.0 |
| Common dandelion | *Taraxacum officinale* | Asteraceae | Asterales | Asia, Europe | | ■ | 0 | 0.0 | 1 | 5.3 | 4.4 | 0.6 | 4.0 | 6.0 |
| White clover | *Trifolium repens* | Fabaceae | Fabales | Asia, Europe | ■ | | 1 | 20.0 | 0 | 0 | 3.1 | 0.2 | 3.0 | 4.0 |

**Table S3.** The 14 pollinator and other flower-visitor insect species recorded through direct observations in *The Lawn is Buzzing* project in Gilpin Park, Merri-Bek (Melbourne, Victoria), their taxonomical associations, the treatment plot they were found at (High: high-intensity mowing; Low: low-intensity mowing), and the number of times they were observed at each treatment. Introduced insect species are indicated with an *.

| Insect group I Species/morphospecies | Family | Mowing intensity | | High-intensity mowing plot | | Low-intensity mowing plot | |
|---|---|---|---|---|---|---|---|
| | | High | Low | Number of records (n=8) | % | Number of records (n=77) | % |
| **Ants** | | | | | | | |
| Formicidae | Formicidae | ■ | ■ | 2 | 25.0 | 11 | 14.3 |
| *Myrmecia pyriformis* | Formicidae | □ | ■ | 0 | 0 | 1 | 1.3 |
| **Bees** | | | | | | | |
| *Apis mellifera* * | Apidae | □ | ■ | 0 | 0 | 14 | 18.2 |
| Lasioglossum | Halictidae | □ | ■ | 0 | 0 | 8 | 10.4 |
| **Beetles** | | | | | | | |
| *Hippodamia variegata* * | Coccinellidae | □ | ■ | 0 | 0 | 2 | 2.6 |
| **Brachyceran flies** | | | | | | | |
| Brachycera | | □ | ■ | 0 | 0 | 6 | 7.8 |
| Syrphidae | Syrphidae | ■ | ■ | 3 | 37.5 | 22 | 28.6 |
| Tephritidae | Tephritidae | □ | ■ | 0 | 0 | 1 | 1.3 |
| Villini | Bombyliidae | □ | ■ | 0 | 0 | 1 | 1.3 |
| **Butterflies** | | | | | | | |
| *Ocybadistes walkeri* | Herperiidae | □ | ■ | 0 | 0 | 2 | 2.6 |
| *Vanessa kershawi* | Nymphalidae | □ | ■ | 0 | 0 | 1 | 1.3 |
| *Zizina otis labradus* | Lycaenidae | ■ | ■ | 3 | 37.5 | 2 | 2.6 |
| **Heteropteran bugs** | | | | | | | |
| *Dindymus versicolor* | Pyrrocoridae | □ | ■ | 0 | 0 | 3 | 3.9 |
| **Parasitoid wasps** | | | | | | | |
| Parasitica | | □ | ■ | 0 | 0 | 3 | 3.9 |

**Table S4.** Name and area of the 15 public parks surveyed within the City of Melbourne (Victoria, Australia) for *The Little Things that Run the City* project, including the park area, the number of surveyed lawn plots in each park and the plot size.

| Park name | Area ($m^2$)* | Plot size ($m^2$) | Number of lawn plots |
|---|---|---|---|
| Royal Park | 1,261,946 | 95 | 9 |
| Princes Park | 329,280 | 95 | 6 |
| Westgate Park | 239,791 | 104 | 4 |
| Fitzroy-Treasury Gardens | 175,776 | 95 | 5 |
| Carlton Gardens South | 47,244 | 106 | 3 |
| University Square | 12,768 | 108 | 2 |
| Lincoln Square | 9,865 | 100 | 2 |
| Argyle Square | 8,335 | 95 | 2 |
| Women's Peace Gardens | 5,684 | 84 | 2 |
| Gardiner Reserve | 3,655 | 148 | 1 |
| Pleasance Gardens | 3,404 | 145 | 1 |
| Murchison Square | 3,294 | 143 | 1 |
| State Library of Victoria | 2,400 | 130 | 1 |
| Canning/Neill Street Reserve | 1,601 | 115 | 1 |
| Garrard Street Reserve | 1,081 | 102 | 1 |

*This figure excludes areas of the park covered by impervious surfaces.

**Table S5.** Estimated species richness for the high-intensity mowing and the low-intensity mowing plots of *The Lawn is Buzzing* project in Gilpin Park, Merri-Bek (Melbourne, Victoria).

| | High-intensity mowing plot | | | | Low-intensity mowing plot | | | |
|---|---|---|---|---|---|---|---|---|
| | **Mean** | **SD** | **2.5%** | **97.5%** | **Mean** | **SD** | **2.5%** | **97.5%** |
| All insects - indigenous | 16.4 | 0.7 | 15.2 | 17.8 | 83.1 | 2.5 | 77.2 | 86.8 |
| Detritivores | 9.1 | 0.7 | 7.8 | 10.4 | 22.5 | 1.6 | 19.2 | 25.2 |
| Herbivores | 5.0 | 0.3 | 4.5 | 5.8 | 21.2 | 0.6 | 19.6 | 22.0 |
| Predators | 2.3 | 0.4 | 2.0 | 3.0 | 3.6 | 0.3 | 3.3 | 4.3 |
| Parasitoids | 2.8 | 0.3 | 2.3 | 3.7 | 19.1 | 1.2 | 16.8 | 21.8 |
| Pollinators | 0.9 | 0.1 | 0.8 | 1.2 | 3.2 | 0.2 | 3.0 | 3.6 |

**Table S6.** Community-level estimates of the probability of occurrence, probability of detection and effect of vegetation height on indigenous insect species for *The Lawn is Buzzing* (Gilpin Park, City of Merri-Bek, Melbourne, Victoria) and *The Little Things that Run the City* (City of Melbourne, Victoria) projects.

| | **Probability of occurrence** | | | | **Probability of detection** | | | | **Effect of vegetation height** | | | |
|---|---|---|---|---|---|---|---|---|---|---|---|---|
| | **Mean** | **SD** | **2.5%** | **97.5%** | **Mean** | **SD** | **2.5%** | **97.5%** | **Mean** | **SD** | **2.5%** | **97.5%** |
| **The Lawn is Buzzing** | | | | | | | | | | | | |
| Whole community | 0.763 | 0.108 | 0.531 | 0.938 | 0.169 | 0.017 | 0.140 | 0.206 | 3.770 | 0.985 | 1.971 | 5.901 |
| Detritivores | 0.655 | 0.150 | 0.385 | 0.937 | 0.226 | 0.056 | 0.136 | 0.353 | 1.762 | 0.869 | 0.670 | 3.983 |
| Herbivores | 0.763 | 0.121 | 0.504 | 0.952 | 0.163 | 0.024 | 0.118 | 0.213 | 4.213 | 1.225 | 2.180 | 6.917 |
| Predators | 0.391 | 0.149 | 0.185 | 0.767 | 0.333 | 0.109 | 0.144 | 0.562 | 0.327 | 0.400 | -0.376 | 1.211 |
| Parasitoids | 0.497 | 0.140 | 0.279 | 0.849 | 0.218 | 0.041 | 0.146 | 0.310 | 1.521 | 0.663 | 0.629 | 3.153 |
| **The Little Things that Run the City** | | | | | | | | | | | | |
| Whole community | 0.341 | 0.085 | 0.199 | 0.531 | 0.027 | 0.005 | 0.019 | 0.038 | 1.249 | 0.322 | 0.718 | 1.959 |

**Table S7.** Species-specific estimates of the probability of occurrence, probability of detection and effect of vegetation height for *The Lawn is Buzzing* project (Gilpin Park, Merri-Bek, Melbourne, Victoria).

| Insect group I Species/morphospecies | Family | Probability of occurrence | | | | Probability of detection | | | | Effect of vegetation height | | | |
|---|---|---|---|---|---|---|---|---|---|---|---|---|---|
| | | Mean | SD | 2.5% | 97.5% | Mean | SD | 2.5% | 97.5% | Mean | SD | 2.5% | 97.5% |
| **Ants [Hymenoptera: Apocrita: Aculeata: Formicoidea]** | | | | | | | | | | | | | |
| *Colobopsis sp. 1* | Formicidae | 0.818 | 0.188 | 0.313 | 0.998 | 0.124 | 0.068 | 0.032 | 0.292 | 3.015 | 1.550 | -0.301 | 5.987 |
| Formicidae 1 | Formicidae | 0.949 | 0.063 | 0.768 | 1.000 | 0.684 | 0.090 | 0.496 | 0.847 | 2.719 | 1.501 | -0.094 | 5.718 |
| Formicidae 2 | Formicidae | 0.849 | 0.153 | 0.429 | 0.998 | 0.157 | 0.074 | 0.049 | 0.334 | 3.189 | 1.522 | -0.091 | 6.120 |
| Formicidae 3 | Formicidae | 0.628 | 0.253 | 0.100 | 0.984 | 0.151 | 0.084 | 0.038 | 0.362 | 3.924 | 1.481 | 1.359 | 7.247 |
| Formicidae 4 | Formicidae | 0.751 | 0.201 | 0.260 | 0.990 | 0.137 | 0.074 | 0.035 | 0.317 | 3.831 | 1.557 | 1.102 | 7.240 |
| Formicidae 5 | Formicidae | 0.754 | 0.199 | 0.260 | 0.992 | 0.136 | 0.074 | 0.034 | 0.315 | 3.847 | 1.536 | 1.092 | 7.226 |
| *Pheidole sp. 1* | Formicidae | 0.909 | 0.107 | 0.604 | 0.999 | 0.290 | 0.095 | 0.130 | 0.496 | 2.720 | 1.729 | -1.281 | 5.783 |
| Ponerinae 1 | Formicidae | 0.697 | 0.218 | 0.207 | 0.985 | 0.143 | 0.077 | 0.038 | 0.332 | 3.981 | 1.542 | 1.284 | 7.425 |
| **Bees [Hymenoptera: Apocrita: Aculeata: Apoidea: Anthophila]** | | | | | | | | | | | | | |
| *Apis mellifera* * | Apidae | 0.604 | 0.268 | 0.078 | 0.985 | 0.266 | 0.118 | 0.086 | 0.540 | 4.014 | 1.444 | 1.535 | 7.271 |
| *Lasioglossum sp. 1* | Halictidae | 0.627 | 0.257 | 0.097 | 0.984 | 0.151 | 0.083 | 0.039 | 0.356 | 3.912 | 1.499 | 1.297 | 7.291 |
| *Lasioglossum sp. 2* | Halictidae | 0.575 | 0.270 | 0.056 | 0.976 | 0.215 | 0.109 | 0.061 | 0.476 | 3.931 | 1.537 | 1.324 | 7.406 |
| *Lasioglossum sp. 3* | Halictidae | 0.822 | 0.147 | 0.445 | 0.991 | 0.416 | 0.119 | 0.203 | 0.660 | 3.822 | 1.464 | 1.197 | 7.034 |
| *Lasioglossum sp. 4* | Halictidae | 0.754 | 0.200 | 0.262 | 0.991 | 0.137 | 0.074 | 0.035 | 0.319 | 3.849 | 1.517 | 1.132 | 7.210 |
| **Beetles [Coleoptera]** | | | | | | | | | | | | | |
| *Anthrenus sp. 1* * | Dermestidae | 0.814 | 0.199 | 0.271 | 0.998 | 0.124 | 0.067 | 0.033 | 0.290 | 3.133 | 1.567 | -0.210 | 6.169 |
| *Hippodamia variegata* * | Coccinellidae | 0.786 | 0.202 | 0.264 | 0.996 | 0.219 | 0.097 | 0.073 | 0.447 | 3.528 | 1.437 | 0.804 | 6.600 |
| *Sitona discoideus* * | Curculionidae | 0.889 | 0.107 | 0.606 | 0.997 | 0.507 | 0.115 | 0.288 | 0.730 | 3.426 | 1.327 | 1.013 | 6.274 |
| Alticini 1 | Chrysomelidae | 0.756 | 0.205 | 0.258 | 0.993 | 0.279 | 0.111 | 0.103 | 0.533 | 3.589 | 1.378 | 1.155 | 6.614 |
| Anthicidae 1 | Anthicidae | 0.632 | 0.255 | 0.098 | 0.983 | 0.152 | 0.084 | 0.037 | 0.359 | 3.931 | 1.524 | 1.272 | 7.346 |
| Anthicidae 2 | Anthicidae | 0.611 | 0.266 | 0.070 | 0.983 | 0.153 | 0.087 | 0.037 | 0.369 | 3.897 | 1.520 | 1.308 | 7.302 |
| Anthicidae 3 | Anthicidae | 0.683 | 0.221 | 0.189 | 0.985 | 0.196 | 0.094 | 0.057 | 0.418 | 4.008 | 1.540 | 1.381 | 7.457 |
| Anthicidae 4 | Anthicidae | 0.694 | 0.222 | 0.188 | 0.986 | 0.144 | 0.078 | 0.037 | 0.332 | 3.956 | 1.537 | 1.293 | 7.381 |
| Anthicidae 5 | Anthicidae | 0.757 | 0.199 | 0.262 | 0.992 | 0.136 | 0.073 | 0.035 | 0.313 | 3.831 | 1.519 | 1.129 | 7.140 |
| Apionini 1 | Brentidae | 0.631 | 0.254 | 0.102 | 0.982 | 0.151 | 0.083 | 0.038 | 0.355 | 3.946 | 1.504 | 1.343 | 7.346 |
| Baridinae 1 | Curculionidae | 0.619 | 0.251 | 0.111 | 0.982 | 0.205 | 0.101 | 0.060 | 0.448 | 3.985 | 1.456 | 1.498 | 7.316 |
| Carabidae 1 | Carabidae | 0.808 | 0.193 | 0.298 | 0.998 | 0.125 | 0.069 | 0.033 | 0.295 | 3.133 | 1.528 | -0.027 | 6.106 |
| Carabidae 2 | Carabidae | 0.634 | 0.252 | 0.111 | 0.984 | 0.150 | 0.082 | 0.039 | 0.353 | 3.959 | 1.519 | 1.391 | 7.399 |
| *Chauliognathus lugubris* | Cantharidea | 0.631 | 0.254 | 0.105 | 0.984 | 0.151 | 0.083 | 0.037 | 0.352 | 3.925 | 1.501 | 1.339 | 7.225 |
| Chrysomelidae 1 | Chrysomelidae | 0.639 | 0.256 | 0.099 | 0.985 | 0.149 | 0.082 | 0.038 | 0.356 | 3.894 | 1.483 | 1.325 | 7.212 |
| *Coccinella transversalis* | Coccinellidae | 0.775 | 0.199 | 0.279 | 0.995 | 0.221 | 0.098 | 0.075 | 0.454 | 3.475 | 1.451 | 0.714 | 6.505 |
| Coccinellidae 1 | Coccinellidae | 0.692 | 0.220 | 0.191 | 0.987 | 0.144 | 0.077 | 0.037 | 0.333 | 3.972 | 1.561 | 1.286 | 7.454 |
| Coccinellidae 2 | Coccinellidae | 0.755 | 0.199 | 0.254 | 0.991 | 0.137 | 0.073 | 0.036 | 0.315 | 3.848 | 1.538 | 1.135 | 7.239 |
| *Cortinicara sp.1* | Latriidae | 0.935 | 0.075 | 0.725 | 0.999 | 0.590 | 0.101 | 0.390 | 0.781 | 2.776 | 1.461 | 0.052 | 5.710 |

Table S7. *Cont.*

| Insect group I Species/morphospecies | Family | Probability of occurrence | | | | Probability of detection | | | | Effect of vegetation height | | | |
|---|---|---|---|---|---|---|---|---|---|---|---|---|---|
| | | Mean | SD | 2.5% | 97.5% | Mean | SD | 2.5% | 97.5% | Mean | SD | 2.5% | 97.5% |
| **Beetles *Cont.*** | | | | | | | | | | | | | |
| Cryptocephalini 1 | Chrysomelidae | 0.755 | 0.197 | 0.263 | 0.991 | 0.137 | 0.073 | 0.036 | 0.316 | 3.836 | 1.546 | 1.057 | 7.252 |
| *Diomus notescens* | Coccinellidae | 0.591 | 0.264 | 0.072 | 0.978 | 0.210 | 0.104 | 0.061 | 0.460 | 3.966 | 1.519 | 1.384 | 7.381 |
| *Diomus sp. 1* | Coccinellidae | 0.621 | 0.267 | 0.069 | 0.986 | 0.153 | 0.086 | 0.039 | 0.365 | 3.930 | 1.516 | 1.332 | 7.365 |
| Elateridae 1 | Elateridae | 0.628 | 0.254 | 0.100 | 0.982 | 0.151 | 0.084 | 0.039 | 0.357 | 3.936 | 1.512 | 1.348 | 7.323 |
| *Euciodes suturalis* | Anthribidae | 0.631 | 0.256 | 0.100 | 0.986 | 0.153 | 0.085 | 0.039 | 0.365 | 3.911 | 1.509 | 1.315 | 7.354 |
| Mordellidae 1 | Mordellidae | 0.621 | 0.257 | 0.089 | 0.982 | 0.152 | 0.085 | 0.037 | 0.363 | 3.941 | 1.500 | 1.339 | 7.320 |
| Mordellidae 2 | Mordellidae | 0.617 | 0.261 | 0.084 | 0.983 | 0.153 | 0.084 | 0.039 | 0.361 | 3.921 | 1.511 | 1.341 | 7.349 |
| Opatrini 1 | Tenebrionidae | 0.811 | 0.190 | 0.305 | 0.998 | 0.126 | 0.070 | 0.032 | 0.299 | 3.157 | 1.531 | -0.015 | 6.150 |
| *Paederus sp. 1* | Staphylinidae | 0.629 | 0.254 | 0.099 | 0.982 | 0.152 | 0.084 | 0.038 | 0.360 | 3.938 | 1.491 | 1.322 | 7.219 |
| Phalacridae 1 | Phalacridae | 0.628 | 0.257 | 0.098 | 0.986 | 0.152 | 0.084 | 0.038 | 0.360 | 3.940 | 1.523 | 1.329 | 7.361 |
| Polyphaga 1 | | 0.626 | 0.258 | 0.094 | 0.983 | 0.151 | 0.084 | 0.038 | 0.356 | 3.947 | 1.582 | 1.289 | 7.413 |
| Polyphaga 2 | | 0.801 | 0.188 | 0.309 | 0.996 | 0.129 | 0.071 | 0.033 | 0.302 | 3.428 | 1.487 | 0.506 | 6.466 |
| *Rhinoncus australis* | Curculionidae | 0.628 | 0.253 | 0.100 | 0.980 | 0.152 | 0.083 | 0.039 | 0.362 | 3.971 | 1.515 | 1.373 | 7.350 |
| **Booklice [Psocoptera]** | | | | | | | | | | | | | |
| Psocoptera 1 | | 0.753 | 0.198 | 0.266 | 0.991 | 0.137 | 0.074 | 0.035 | 0.319 | 3.814 | 1.522 | 1.090 | 7.123 |
| **Brachyceran flies [Diptera: Brachycera]** | | | | | | | | | | | | | |
| *Anabarhynchus sp. 1* | Therevidae | 0.627 | 0.258 | 0.096 | 0.984 | 0.152 | 0.085 | 0.038 | 0.359 | 3.907 | 1.495 | 1.281 | 7.230 |
| *Anthomyia punctipennis* | Anthomyiidae | 0.760 | 0.200 | 0.276 | 0.994 | 0.277 | 0.111 | 0.104 | 0.534 | 3.599 | 1.373 | 1.155 | 6.633 |
| Anthomyiidae 1 | Anthomyiidae | 0.795 | 0.189 | 0.313 | 0.996 | 0.171 | 0.083 | 0.051 | 0.369 | 3.522 | 1.427 | 0.904 | 6.632 |
| Anthomyiidae 2 | Anthomyiidae | 0.864 | 0.129 | 0.527 | 0.998 | 0.466 | 0.120 | 0.246 | 0.709 | 3.179 | 1.356 | 0.719 | 6.014 |
| *Atherigona sp. 1* | Muscidae | 0.684 | 0.220 | 0.191 | 0.984 | 0.194 | 0.093 | 0.058 | 0.413 | 3.993 | 1.517 | 1.364 | 7.382 |
| *Australoactina sp. 1* | Stratiomyidae | 0.748 | 0.196 | 0.272 | 0.988 | 0.182 | 0.087 | 0.055 | 0.391 | 3.873 | 1.510 | 1.236 | 7.248 |
| *Austrotephritis poenia* | Tephritidae | 0.771 | 0.203 | 0.267 | 0.994 | 0.224 | 0.101 | 0.074 | 0.462 | 3.510 | 1.409 | 0.862 | 6.445 |
| Brachycera 1 | | 0.797 | 0.168 | 0.388 | 0.994 | 0.531 | 0.129 | 0.287 | 0.785 | 3.527 | 1.328 | 1.239 | 6.494 |
| Brachycera 2 | | 0.632 | 0.257 | 0.094 | 0.985 | 0.151 | 0.083 | 0.038 | 0.354 | 3.901 | 1.488 | 1.288 | 7.229 |
| Brachycera 3 | | 0.778 | 0.195 | 0.298 | 0.996 | 0.267 | 0.109 | 0.099 | 0.521 | 3.334 | 1.358 | 0.826 | 6.176 |
| Brachycera 4 | | 0.810 | 0.185 | 0.325 | 0.998 | 0.166 | 0.081 | 0.050 | 0.360 | 3.227 | 1.439 | 0.424 | 6.175 |
| Brachycera 5 | | 0.805 | 0.189 | 0.313 | 0.998 | 0.167 | 0.082 | 0.050 | 0.364 | 3.267 | 1.397 | 0.608 | 6.163 |
| Brachycera 6 | | 0.810 | 0.194 | 0.291 | 0.998 | 0.126 | 0.069 | 0.033 | 0.298 | 3.147 | 1.513 | 0.007 | 6.064 |
| Brachycera 7 | | 0.893 | 0.115 | 0.571 | 0.998 | 0.224 | 0.087 | 0.087 | 0.420 | 3.059 | 1.572 | -0.347 | 6.076 |
| Brachycera 8 | | 0.603 | 0.262 | 0.086 | 0.982 | 0.209 | 0.104 | 0.060 | 0.462 | 3.930 | 1.520 | 1.278 | 7.294 |
| Brachycera 9 | | 0.634 | 0.252 | 0.109 | 0.983 | 0.152 | 0.085 | 0.038 | 0.361 | 3.944 | 1.516 | 1.339 | 7.326 |
| Brachycera 10 | | 0.697 | 0.217 | 0.200 | 0.982 | 0.303 | 0.120 | 0.113 | 0.573 | 3.911 | 1.528 | 1.269 | 7.345 |
| Brachycera 11 | | 0.614 | 0.260 | 0.087 | 0.980 | 0.153 | 0.085 | 0.039 | 0.365 | 3.921 | 1.491 | 1.325 | 7.275 |
| Brachycera 12 | | 0.792 | 0.190 | 0.295 | 0.995 | 0.132 | 0.071 | 0.034 | 0.308 | 3.539 | 1.516 | 0.590 | 6.735 |
| Brachycera 13 | | 0.622 | 0.261 | 0.081 | 0.982 | 0.153 | 0.086 | 0.039 | 0.365 | 3.915 | 1.487 | 1.319 | 7.234 |

**Table S7.** ***Cont.***

| Insect group l Species/morphospecies | Family | Probability of occurrence | | | | Probability of detection | | | | Effect of vegetation height | | | |
|---|---|---|---|---|---|---|---|---|---|---|---|---|---|
| | | Mean | SD | 2.5% | 97.5% | Mean | SD | 2.5% | 97.5% | Mean | SD | 2.5% | 97.5% |
| **Brachyceran flies *Cont.*** | | | | | | | | | | | | | |
| *Cerdistus sp. 1* | Asilidae | 0.617 | 0.265 | 0.076 | 0.983 | 0.154 | 0.088 | 0.038 | 0.370 | 3.869 | 1.474 | 1.331 | 7.145 |
| *Coenosia sp. 1* | Muscidae | 0.911 | 0.089 | 0.670 | 0.998 | 0.564 | 0.108 | 0.349 | 0.771 | 3.341 | 1.350 | 0.839 | 6.195 |
| *Depressa atrata* | Lauxaniidae | 0.626 | 0.257 | 0.097 | 0.984 | 0.152 | 0.083 | 0.039 | 0.357 | 3.951 | 1.508 | 1.394 | 7.368 |
| Diaphorinae 1 | Dolichopodidae | 0.635 | 0.254 | 0.106 | 0.983 | 0.150 | 0.082 | 0.038 | 0.352 | 3.921 | 1.486 | 1.358 | 7.205 |
| *Dichetophora sp. 1* | Sciomyzidae | 0.574 | 0.256 | 0.087 | 0.973 | 0.271 | 0.117 | 0.090 | 0.541 | 4.035 | 1.497 | 1.526 | 7.402 |
| Dolichopodidae 1 | Dolichopodidae | 0.625 | 0.254 | 0.096 | 0.982 | 0.151 | 0.082 | 0.039 | 0.353 | 3.914 | 1.496 | 1.304 | 7.209 |
| Hybotidae 1 | Hybotidae | 0.619 | 0.259 | 0.088 | 0.982 | 0.153 | 0.084 | 0.038 | 0.358 | 3.908 | 1.502 | 1.285 | 7.236 |
| *Hydrellia tritici* | Ephydridae | 0.949 | 0.062 | 0.775 | 1.000 | 0.756 | 0.083 | 0.578 | 0.899 | 2.741 | 1.496 | -0.058 | 5.735 |
| Muscidae 1 | Muscidae | 0.666 | 0.221 | 0.189 | 0.978 | 0.252 | 0.108 | 0.084 | 0.500 | 4.044 | 1.486 | 1.439 | 7.366 |
| Muscidae 2 | Muscidae | 0.859 | 0.136 | 0.494 | 0.998 | 0.196 | 0.084 | 0.067 | 0.391 | 3.292 | 1.481 | 0.302 | 6.246 |
| Oestroidea 1 | | 0.791 | 0.189 | 0.308 | 0.995 | 0.172 | 0.083 | 0.053 | 0.372 | 3.514 | 1.441 | 0.816 | 6.603 |
| Oestroidea 2 | | 0.627 | 0.254 | 0.103 | 0.986 | 0.152 | 0.084 | 0.038 | 0.359 | 3.949 | 1.499 | 1.342 | 7.318 |
| Oestroidea 3 | | 0.817 | 0.189 | 0.306 | 0.999 | 0.124 | 0.068 | 0.032 | 0.290 | 3.008 | 1.554 | -0.288 | 5.955 |
| Orthorrhapha 1 | | 0.800 | 0.189 | 0.300 | 0.996 | 0.128 | 0.071 | 0.033 | 0.301 | 3.399 | 1.492 | 0.410 | 6.470 |
| *Parapalaeosepsis plebeia* | Sepsidae | 0.815 | 0.139 | 0.472 | 0.985 | 0.693 | 0.113 | 0.452 | 0.885 | 3.916 | 1.307 | 1.630 | 6.847 |
| *Phasia sp. 1* | Tachinidae | 0.621 | 0.264 | 0.078 | 0.982 | 0.151 | 0.084 | 0.038 | 0.357 | 3.892 | 1.495 | 1.275 | 7.278 |
| *Poecilohetaerus aquilus* | Lauxaniidae | 0.910 | 0.093 | 0.654 | 0.998 | 0.520 | 0.109 | 0.310 | 0.733 | 3.264 | 1.414 | 0.514 | 6.153 |
| *Rivellia sp. 1* | Platystomatidae | 0.620 | 0.264 | 0.085 | 0.984 | 0.152 | 0.084 | 0.038 | 0.362 | 3.890 | 1.486 | 1.264 | 7.213 |
| *Sapromyza mallochiana* | Lauxaniidae | 0.545 | 0.257 | 0.070 | 0.964 | 0.349 | 0.135 | 0.127 | 0.643 | 4.013 | 1.447 | 1.539 | 7.311 |
| *Simosyrphus grandicornis* | Syrphidae | 0.809 | 0.186 | 0.325 | 0.998 | 0.166 | 0.080 | 0.049 | 0.357 | 3.266 | 1.425 | 0.498 | 6.203 |
| *Spathulina acroleuca* | Tephritidae | 0.793 | 0.190 | 0.300 | 0.995 | 0.170 | 0.082 | 0.051 | 0.370 | 3.498 | 1.433 | 0.795 | 6.530 |
| *Sphenella ruficeps* | Tephritidae | 0.746 | 0.197 | 0.274 | 0.990 | 0.182 | 0.087 | 0.054 | 0.390 | 3.875 | 1.492 | 1.286 | 7.220 |
| *Steganopsis melanogaster* | Lauxaniidae | 0.793 | 0.172 | 0.366 | 0.992 | 0.430 | 0.127 | 0.205 | 0.693 | 3.640 | 1.366 | 1.184 | 6.631 |
| Therevidae 1 | Therevidae | 0.623 | 0.253 | 0.103 | 0.981 | 0.153 | 0.085 | 0.039 | 0.362 | 3.936 | 1.509 | 1.328 | 7.259 |
| **Earwigs [Dermaptera]** | | | | | | | | | | | | | |
| *Forficula dentata* * | Forficulidae | 0.599 | 0.272 | 0.073 | 0.985 | 0.268 | 0.118 | 0.088 | 0.540 | 4.035 | 1.499 | 1.512 | 7.373 |
| **Grasshoppers, crickets and katydids [Orthoptera]** | | | | | | | | | | | | | |
| Catantopini 1 | Acrididae | 0.694 | 0.221 | 0.194 | 0.987 | 0.143 | 0.077 | 0.036 | 0.328 | 3.965 | 1.530 | 1.330 | 7.355 |
| *Conocephalus albescens* | Tettigoniidae | 0.625 | 0.259 | 0.090 | 0.985 | 0.152 | 0.084 | 0.039 | 0.358 | 3.896 | 1.482 | 1.331 | 7.233 |
| *Conocephalus upoluensis* | Tettigoniidae | 0.626 | 0.258 | 0.091 | 0.983 | 0.153 | 0.084 | 0.038 | 0.359 | 3.895 | 1.499 | 1.318 | 7.215 |
| Gryllidae 1 | Gryllidae | 0.621 | 0.265 | 0.084 | 0.986 | 0.153 | 0.085 | 0.038 | 0.363 | 3.904 | 1.523 | 1.306 | 7.246 |
| *Schizobothrus flavovittatus* | Acrididae | 0.740 | 0.203 | 0.250 | 0.991 | 0.184 | 0.090 | 0.054 | 0.397 | 3.877 | 1.509 | 1.235 | 7.238 |
| Trigonidiinae 1 | Trigonidiidae | 0.585 | 0.271 | 0.059 | 0.978 | 0.213 | 0.109 | 0.061 | 0.480 | 3.920 | 1.512 | 1.311 | 7.333 |

**Table S7.** ***Cont.***

| Insect group I Species/morphospecies | Family | Probability of occurrence | | | | Probability of detection | | | | Effect of vegetation height | | | |
|---|---|---|---|---|---|---|---|---|---|---|---|---|---|
| | | Mean | SD | 2.5% | 97.5% | Mean | SD | 2.5% | 97.5% | Mean | SD | 2.5% | 97.5% |
| **Heteropteran bugs [Hemiptera: Heteroptera]** | | | | | | | | | | | | | |
| *Pyrrhocoris apterus ** | Pyrrocoridae | 0.781 | 0.177 | 0.340 | 0.992 | 0.225 | 0.095 | 0.078 | 0.444 | 4.005 | 1.495 | 1.450 | 7.339 |
| *Amorbus sp. 1* | Coreidae | 0.611 | 0.267 | 0.076 | 0.985 | 0.154 | 0.086 | 0.039 | 0.369 | 3.902 | 1.503 | 1.307 | 7.269 |
| *Cermatulus nasalis* | Pentatomidae | 0.625 | 0.255 | 0.100 | 0.982 | 0.151 | 0.084 | 0.039 | 0.359 | 3.911 | 1.470 | 1.330 | 7.177 |
| Cydnidae 1 | Cydnidae | 0.698 | 0.219 | 0.194 | 0.985 | 0.143 | 0.077 | 0.036 | 0.329 | 3.953 | 1.552 | 1.223 | 7.355 |
| *Dindymus versicolor* | Pyrrocoridae | 0.611 | 0.259 | 0.088 | 0.979 | 0.209 | 0.103 | 0.060 | 0.453 | 3.967 | 1.536 | 1.300 | 7.404 |
| *Eribotes sp. 2* | Pentatomidae | 0.756 | 0.198 | 0.262 | 0.992 | 0.136 | 0.073 | 0.036 | 0.317 | 3.813 | 1.556 | 1.035 | 7.228 |
| *Melanacanthus sp. 1* | Alydidae | 0.615 | 0.264 | 0.084 | 0.985 | 0.153 | 0.086 | 0.038 | 0.365 | 3.913 | 1.489 | 1.384 | 7.292 |
| Miridae 1 | Miridae | 0.621 | 0.261 | 0.085 | 0.984 | 0.154 | 0.085 | 0.039 | 0.361 | 3.914 | 1.507 | 1.341 | 7.305 |
| Miridae 2 | Miridae | 0.753 | 0.201 | 0.257 | 0.991 | 0.138 | 0.075 | 0.035 | 0.325 | 3.832 | 1.538 | 1.024 | 7.172 |
| *Mutsca brevicornis* | Alydidae | 0.633 | 0.256 | 0.103 | 0.985 | 0.149 | 0.083 | 0.037 | 0.352 | 3.953 | 1.504 | 1.384 | 7.356 |
| *Nabis kinbergii* | Nabidae | 0.634 | 0.218 | 0.176 | 0.964 | 0.383 | 0.129 | 0.161 | 0.652 | 4.160 | 1.487 | 1.674 | 7.537 |
| *Nysius vinitor* | Lygaeidae | 0.801 | 0.170 | 0.378 | 0.995 | 0.263 | 0.102 | 0.100 | 0.491 | 3.640 | 1.369 | 1.177 | 6.641 |
| *Oxycarenus luctuosus* | Oxycarenidae | 0.620 | 0.258 | 0.093 | 0.982 | 0.153 | 0.085 | 0.037 | 0.361 | 3.937 | 1.494 | 1.365 | 7.295 |
| Pentatomidae 1 | Pentatomidae | 0.751 | 0.200 | 0.259 | 0.991 | 0.137 | 0.073 | 0.035 | 0.316 | 3.804 | 1.531 | 0.963 | 7.085 |
| *Sidnia kinbergi* | Miridae | 0.619 | 0.257 | 0.087 | 0.981 | 0.153 | 0.084 | 0.039 | 0.357 | 3.940 | 1.501 | 1.362 | 7.258 |
| *Stenophylla macreta* | Pachygronthidae | 0.563 | 0.261 | 0.071 | 0.967 | 0.279 | 0.122 | 0.091 | 0.561 | 3.993 | 1.469 | 1.526 | 7.387 |
| Thaumastocoridae 1 | Thaumastocoridae | 0.699 | 0.220 | 0.201 | 0.989 | 0.144 | 0.077 | 0.038 | 0.331 | 3.943 | 1.534 | 1.229 | 7.342 |
| **Jumping plant lice [Hemiptera: Sternorrhyncha: Psylloidea]** | | | | | | | | | | | | | |
| Psylloidea 1 | | 0.634 | 0.256 | 0.101 | 0.986 | 0.150 | 0.083 | 0.038 | 0.352 | 3.920 | 1.540 | 1.281 | 7.433 |
| Psylloidea 2 | | 0.631 | 0.255 | 0.104 | 0.983 | 0.151 | 0.082 | 0.039 | 0.353 | 3.949 | 1.523 | 1.358 | 7.371 |
| Psylloidea 3 | | 0.810 | 0.190 | 0.303 | 0.998 | 0.126 | 0.069 | 0.033 | 0.298 | 3.148 | 1.526 | -0.029 | 6.153 |
| **Lacewings [Neuroptera]** | | | | | | | | | | | | | |
| *Micromus tasmaniae* | Hemerobiidae | 0.697 | 0.206 | 0.232 | 0.979 | 0.362 | 0.124 | 0.152 | 0.628 | 4.017 | 1.473 | 1.509 | 7.351 |
| **Leafhoppers [Hemiptera: Auchenorrhyncha: Membracoidea]** | | | | | | | | | | | | | |
| *Brunotartessus fulvus* | Cicadellidae | 0.798 | 0.192 | 0.295 | 0.996 | 0.130 | 0.070 | 0.034 | 0.301 | 3.446 | 1.479 | 0.561 | 6.533 |
| Cicadellidae 1 | Cicadellidae | 0.835 | 0.159 | 0.411 | 0.996 | 0.297 | 0.108 | 0.124 | 0.544 | 3.411 | 1.465 | 0.578 | 6.433 |
| Cicadellidae 2 | Cicadellidae | 0.613 | 0.253 | 0.100 | 0.980 | 0.206 | 0.101 | 0.060 | 0.446 | 4.009 | 1.481 | 1.501 | 7.426 |
| Cicadellidae 3 | Cicadellidae | 0.813 | 0.191 | 0.303 | 0.998 | 0.125 | 0.067 | 0.033 | 0.290 | 3.135 | 1.521 | -0.038 | 6.120 |
| Cicadellidae 4 | Cicadellidae | 0.792 | 0.197 | 0.293 | 0.998 | 0.214 | 0.097 | 0.072 | 0.444 | 3.266 | 1.438 | 0.465 | 6.187 |
| Cicadellidae 5 | Cicadellidae | 0.852 | 0.135 | 0.500 | 0.997 | 0.288 | 0.101 | 0.121 | 0.508 | 3.416 | 1.356 | 0.860 | 6.311 |
| Cicadellidae 6 | Cicadellidae | 0.631 | 0.253 | 0.099 | 0.984 | 0.151 | 0.083 | 0.037 | 0.357 | 3.891 | 1.512 | 1.231 | 7.204 |
| Cicadellidae 7 | Cicadellidae | 0.789 | 0.176 | 0.346 | 0.992 | 0.269 | 0.104 | 0.103 | 0.503 | 3.750 | 1.444 | 1.176 | 6.928 |
| Cicadellidae 8 | Cicadellidae | 0.679 | 0.222 | 0.194 | 0.985 | 0.194 | 0.092 | 0.058 | 0.410 | 4.046 | 1.526 | 1.444 | 7.458 |
| Cicadellidae 9 | Cicadellidae | 0.820 | 0.167 | 0.388 | 0.997 | 0.207 | 0.090 | 0.070 | 0.419 | 3.325 | 1.406 | 0.623 | 6.209 |

**Table S7. *Cont.***

| Insect group I Species/morphospecies | Family | Probability of occurrence | | | | Probability of detection | | | | Effect of vegetation height | | | |
|---|---|---|---|---|---|---|---|---|---|---|---|---|---|
| | | Mean | SD | 2.5% | 97.5% | Mean | SD | 2.5% | 97.5% | Mean | SD | 2.5% | 97.5% |
| **Leafhoppers *Cont.*** | | | | | | | | | | | | | |
| Cicadellidae 10 | Cicadellidae | 0.697 | 0.219 | 0.201 | 0.987 | 0.144 | 0.078 | 0.037 | 0.334 | 3.939 | 1.524 | 1.274 | 7.367 |
| Cicadellidae 11 | Cicadellidae | 0.695 | 0.222 | 0.186 | 0.985 | 0.144 | 0.077 | 0.037 | 0.334 | 3.954 | 1.521 | 1.310 | 7.365 |
| Cicadellidae 12 | Cicadellidae | 0.701 | 0.219 | 0.195 | 0.987 | 0.143 | 0.078 | 0.037 | 0.336 | 3.945 | 1.541 | 1.267 | 7.407 |
| Cicadellidae 13 | Cicadellidae | 0.697 | 0.221 | 0.195 | 0.988 | 0.143 | 0.077 | 0.037 | 0.331 | 3.946 | 1.531 | 1.220 | 7.287 |
| Cicadellidae 14 | Cicadellidae | 0.748 | 0.201 | 0.249 | 0.989 | 0.138 | 0.074 | 0.036 | 0.320 | 3.833 | 1.537 | 1.082 | 7.171 |
| Cicadellidae 15 | Cicadellidae | 0.753 | 0.199 | 0.262 | 0.992 | 0.137 | 0.074 | 0.035 | 0.316 | 3.817 | 1.547 | 1.070 | 7.211 |
| **Nematoceran flies [Diptera: Nematocera]** | | | | | | | | | | | | | |
| Tipulomorpha 1 | | 0.695 | 0.221 | 0.202 | 0.987 | 0.144 | 0.078 | 0.037 | 0.335 | 3.950 | 1.521 | 1.260 | 7.386 |
| **Parasitoid wasps [Hymenoptera: Apocrita: Parasitica]** | | | | | | | | | | | | | |
| Alysiinae 1 | Braconidae | 0.751 | 0.201 | 0.257 | 0.992 | 0.137 | 0.074 | 0.035 | 0.319 | 3.785 | 1.558 | 0.945 | 7.187 |
| Braconidae 1 | Braconidae | 0.615 | 0.252 | 0.103 | 0.977 | 0.206 | 0.100 | 0.060 | 0.445 | 3.985 | 1.487 | 1.464 | 7.388 |
| Braconidae 2 | Braconidae | 0.850 | 0.151 | 0.446 | 0.998 | 0.197 | 0.085 | 0.067 | 0.394 | 3.153 | 1.455 | 0.286 | 6.052 |
| Braconidae 3 | Braconidae | 0.694 | 0.224 | 0.191 | 0.986 | 0.143 | 0.077 | 0.037 | 0.332 | 3.935 | 1.547 | 1.214 | 7.363 |
| Braconidae 4 | Braconidae | 0.778 | 0.173 | 0.354 | 0.989 | 0.177 | 0.084 | 0.053 | 0.378 | 3.911 | 1.519 | 1.258 | 7.208 |
| Braconidae 5 | Braconidae | 0.852 | 0.156 | 0.427 | 0.999 | 0.157 | 0.076 | 0.047 | 0.340 | 3.045 | 1.557 | -0.262 | 5.999 |
| Campopleginae 1 | Ichneumonidae | 0.740 | 0.205 | 0.248 | 0.991 | 0.183 | 0.088 | 0.055 | 0.392 | 3.875 | 1.500 | 1.273 | 7.218 |
| Chalcididae 1 | Chalcididae | 0.694 | 0.220 | 0.200 | 0.985 | 0.144 | 0.078 | 0.037 | 0.336 | 3.938 | 1.516 | 1.271 | 7.324 |
| Chalcidoidea 1 | | 0.623 | 0.262 | 0.085 | 0.984 | 0.152 | 0.085 | 0.037 | 0.363 | 3.925 | 1.503 | 1.343 | 7.278 |
| Chalcidoidea 2 | | 0.615 | 0.262 | 0.077 | 0.982 | 0.153 | 0.086 | 0.038 | 0.367 | 3.921 | 1.495 | 1.338 | 7.241 |
| Chalcidoidea 3 | | 0.776 | 0.175 | 0.342 | 0.990 | 0.177 | 0.084 | 0.053 | 0.378 | 3.912 | 1.511 | 1.228 | 7.206 |
| Cheloninae 1 | Braconidae | 0.595 | 0.263 | 0.078 | 0.979 | 0.211 | 0.104 | 0.062 | 0.462 | 3.940 | 1.484 | 1.356 | 7.281 |
| Cheloninae 2 | Braconidae | 0.780 | 0.193 | 0.295 | 0.994 | 0.175 | 0.086 | 0.052 | 0.380 | 3.616 | 1.441 | 0.982 | 6.746 |
| Cheloninae 3 | Braconidae | 0.755 | 0.199 | 0.268 | 0.991 | 0.137 | 0.074 | 0.035 | 0.318 | 3.823 | 1.495 | 1.163 | 7.082 |
| *Enicospilus sp. 1* | Ichneumonidae | 0.624 | 0.258 | 0.093 | 0.985 | 0.152 | 0.083 | 0.039 | 0.357 | 3.937 | 1.501 | 1.359 | 7.340 |
| *Epitranus sp. 1* | Chalcididae | 0.696 | 0.217 | 0.204 | 0.987 | 0.143 | 0.078 | 0.036 | 0.334 | 3.965 | 1.524 | 1.271 | 7.305 |
| *Gasteruption raphidioides* | Gasteruptiidae | 0.699 | 0.219 | 0.205 | 0.989 | 0.142 | 0.078 | 0.036 | 0.332 | 3.926 | 1.531 | 1.245 | 7.353 |
| Gravenhorstiini 1 | Ichneumonidae | 0.629 | 0.260 | 0.087 | 0.983 | 0.152 | 0.085 | 0.037 | 0.360 | 3.903 | 1.497 | 1.268 | 7.213 |
| Ichneumonoidea 1 | | 0.766 | 0.179 | 0.332 | 0.991 | 0.228 | 0.097 | 0.078 | 0.455 | 3.939 | 1.496 | 1.260 | 7.241 |
| *Lissonota sp. 1* | Ichneumonidae | 0.734 | 0.210 | 0.229 | 0.988 | 0.185 | 0.090 | 0.055 | 0.397 | 3.812 | 1.584 | 0.923 | 7.246 |
| *Myrmeleonostenus sp. 1* | Ichneumonidae | 0.750 | 0.198 | 0.261 | 0.990 | 0.182 | 0.087 | 0.054 | 0.391 | 3.921 | 1.532 | 1.294 | 7.340 |

**Table S7.** ***Cont.***

| Insect group I Species/morphospecies | Family | Probability of occurrence | | | | Probability of detection | | | | Effect of vegetation height | | | |
|---|---|---|---|---|---|---|---|---|---|---|---|---|---|
| | | Mean | SD | 2.5% | 97.5% | Mean | SD | 2.5% | 97.5% | Mean | SD | 2.5% | 97.5% |
| **Parasitoid wasps *Cont.*** | | | | | | | | | | | | | |
| Parasitica 1 | | 0.803 | 0.188 | 0.308 | 0.996 | 0.129 | 0.070 | 0.033 | 0.299 | 3.437 | 1.507 | 0.454 | 6.497 |
| Parasitica 2 | | 0.626 | 0.256 | 0.095 | 0.982 | 0.152 | 0.083 | 0.038 | 0.361 | 3.941 | 1.498 | 1.329 | 7.291 |
| Parasitica 3 | | 0.665 | 0.219 | 0.192 | 0.980 | 0.252 | 0.108 | 0.084 | 0.497 | 4.067 | 1.469 | 1.564 | 7.388 |
| Parasitica 4 | | 0.567 | 0.263 | 0.071 | 0.974 | 0.278 | 0.124 | 0.089 | 0.564 | 3.987 | 1.490 | 1.469 | 7.342 |
| Parasitica 5 | | 0.626 | 0.256 | 0.095 | 0.981 | 0.152 | 0.084 | 0.038 | 0.358 | 3.922 | 1.508 | 1.322 | 7.282 |
| Parasitica 6 | | 0.605 | 0.259 | 0.086 | 0.979 | 0.208 | 0.105 | 0.060 | 0.464 | 3.958 | 1.511 | 1.350 | 7.342 |
| Parasitica 7 | | 0.631 | 0.255 | 0.095 | 0.982 | 0.151 | 0.083 | 0.037 | 0.356 | 3.935 | 1.499 | 1.339 | 7.307 |
| Parasitica 8 | | 0.634 | 0.251 | 0.114 | 0.982 | 0.150 | 0.082 | 0.038 | 0.352 | 3.912 | 1.532 | 1.262 | 7.359 |
| Parasitica 9 | | 0.633 | 0.252 | 0.108 | 0.982 | 0.152 | 0.083 | 0.038 | 0.357 | 3.916 | 1.498 | 1.335 | 7.267 |
| Parasitica 10 | | 0.796 | 0.168 | 0.377 | 0.992 | 0.268 | 0.104 | 0.102 | 0.503 | 3.799 | 1.392 | 1.307 | 6.836 |
| Parasitica 11 | | 0.625 | 0.258 | 0.088 | 0.983 | 0.153 | 0.084 | 0.039 | 0.358 | 3.902 | 1.466 | 1.351 | 7.167 |
| Parasitica 12 | | 0.569 | 0.259 | 0.080 | 0.971 | 0.276 | 0.121 | 0.092 | 0.551 | 3.993 | 1.450 | 1.545 | 7.267 |
| Parasitica 13 | | 0.762 | 0.206 | 0.253 | 0.994 | 0.225 | 0.100 | 0.075 | 0.462 | 3.623 | 1.457 | 0.960 | 6.750 |
| Parasitica 14 | | 0.580 | 0.268 | 0.068 | 0.977 | 0.215 | 0.110 | 0.061 | 0.482 | 3.951 | 1.524 | 1.389 | 7.435 |
| Parasitica 15 | | 0.618 | 0.261 | 0.084 | 0.983 | 0.153 | 0.085 | 0.039 | 0.366 | 3.923 | 1.500 | 1.360 | 7.304 |
| Parasitica 16 | | 0.776 | 0.175 | 0.339 | 0.990 | 0.178 | 0.084 | 0.055 | 0.378 | 3.974 | 1.593 | 1.228 | 7.528 |
| Parasitica 17 | | 0.696 | 0.219 | 0.202 | 0.985 | 0.145 | 0.079 | 0.037 | 0.336 | 3.933 | 1.527 | 1.260 | 7.268 |
| Parasitica 18 | | 0.775 | 0.175 | 0.343 | 0.988 | 0.178 | 0.084 | 0.054 | 0.375 | 3.916 | 1.504 | 1.269 | 7.173 |
| Parasitica 19 | | 0.699 | 0.219 | 0.199 | 0.987 | 0.143 | 0.077 | 0.037 | 0.330 | 3.972 | 1.559 | 1.282 | 7.456 |
| Parasitica 20 | | 0.696 | 0.220 | 0.196 | 0.986 | 0.145 | 0.077 | 0.038 | 0.332 | 3.950 | 1.526 | 1.280 | 7.317 |
| Parasitica 21 | | 0.694 | 0.221 | 0.196 | 0.988 | 0.144 | 0.079 | 0.036 | 0.336 | 3.944 | 1.526 | 1.292 | 7.330 |
| Parasitica 22 | | 0.693 | 0.222 | 0.190 | 0.988 | 0.144 | 0.078 | 0.037 | 0.338 | 3.969 | 1.522 | 1.276 | 7.313 |
| Parasitica 23 | | 0.751 | 0.201 | 0.265 | 0.992 | 0.136 | 0.073 | 0.035 | 0.313 | 3.797 | 1.555 | 1.023 | 7.163 |
| Parasitica 24 | | 0.754 | 0.200 | 0.258 | 0.993 | 0.136 | 0.074 | 0.036 | 0.318 | 3.802 | 1.555 | 0.968 | 7.143 |
| Parasitica 25 | | 0.747 | 0.202 | 0.255 | 0.990 | 0.138 | 0.074 | 0.036 | 0.320 | 3.799 | 1.516 | 1.072 | 7.105 |
| *Setanta sp. 1* | Ichneumonidae | 0.630 | 0.254 | 0.097 | 0.983 | 0.151 | 0.083 | 0.038 | 0.355 | 3.937 | 1.537 | 1.278 | 7.364 |
| Tersilochinae 1 | Ichneumonidae | 0.622 | 0.255 | 0.093 | 0.980 | 0.152 | 0.085 | 0.038 | 0.362 | 3.938 | 1.512 | 1.372 | 7.319 |
| **Planthoppers [Hemiptera: Auchenorryncha: Fulgoromorpha]** | | | | | | | | | | | | | |
| *Anzora unicolor* | Flatidae | 0.692 | 0.222 | 0.188 | 0.986 | 0.144 | 0.078 | 0.036 | 0.335 | 3.918 | 1.573 | 1.192 | 7.313 |
| Delphacidae 1 | Delphacidae | 0.711 | 0.209 | 0.233 | 0.984 | 0.298 | 0.116 | 0.113 | 0.558 | 3.984 | 1.482 | 1.461 | 7.292 |
| Delphacidae 2 | Delphacidae | 0.631 | 0.259 | 0.093 | 0.984 | 0.151 | 0.083 | 0.038 | 0.355 | 3.962 | 1.507 | 1.379 | 7.386 |
| Delphacidae 3 | Delphacidae | 0.757 | 0.197 | 0.268 | 0.992 | 0.137 | 0.074 | 0.035 | 0.316 | 3.864 | 1.508 | 1.157 | 7.156 |

**Table S7.** ***Cont.***

| Insect group I Species/morphospecies | Family | Probability of occurrence | | | | Probability of detection | | | | Effect of vegetation height | | | |
|---|---|---|---|---|---|---|---|---|---|---|---|---|---|
| | | Mean | SD | 2.5% | 97.5% | Mean | SD | 2.5% | 97.5% | Mean | SD | 2.5% | 97.5% |
| **Stinging wasps [Hymenoptera: Apocrita: Aculeata, excl. Apoidea and Formicoidea]** | | | | | | | | | | | | | |
| Bethylidae 1 | Bethylidae | 0.627 | 0.257 | 0.099 | 0.986 | 0.151 | 0.083 | 0.037 | 0.357 | 3.913 | 1.509 | 1.292 | 7.308 |
| Bethylidae 2 | Bethylidae | 0.615 | 0.265 | 0.082 | 0.983 | 0.154 | 0.087 | 0.037 | 0.373 | 3.893 | 1.488 | 1.353 | 7.253 |
| Bethylidae 3 | Bethylidae | 0.837 | 0.140 | 0.478 | 0.993 | 0.303 | 0.106 | 0.125 | 0.536 | 3.754 | 1.437 | 1.124 | 6.867 |
| Dasymutillini 1 | Mutillidae | 0.707 | 0.216 | 0.213 | 0.988 | 0.143 | 0.077 | 0.037 | 0.330 | 3.933 | 1.558 | 1.168 | 7.392 |
| Pompilidae 1 | Pompilidae | 0.625 | 0.258 | 0.092 | 0.984 | 0.153 | 0.085 | 0.037 | 0.365 | 3.948 | 1.497 | 1.390 | 7.330 |
| Pompilidae 2 | Pompilidae | 0.617 | 0.260 | 0.084 | 0.982 | 0.153 | 0.086 | 0.039 | 0.365 | 3.896 | 1.488 | 1.316 | 7.201 |
| Thynnidae 1 | Thynnidae | 0.697 | 0.222 | 0.190 | 0.989 | 0.144 | 0.078 | 0.036 | 0.336 | 3.940 | 1.512 | 1.245 | 7.278 |

**Table S8.** Estimated insect species richness for the lawns surveyed within the City of Melbourne (Victoria, Australia) for *The Little Things that Run the City* (TLT) and *The Lawn is Buzzing* (TLB) projects.

| Lawn | Project | Species richness | | | |
|---|---|---|---|---|---|
| | | Mean | SD | 2.5% | 97.5% |
| Low intensity mowing | TLB | 83.1 | 2.5 | 77.2 | 86.8 |
| Royal Park 1 | TLT | 35.4 | 2.2 | 30.0 | 38.0 |
| Fitzroy-Treasury Gardens 2 | TLT | 34.9 | 2.4 | 29.0 | 38.0 |
| Carlton Gardens South 3 | TLT | 31.6 | 1.6 | 28.0 | 33.0 |
| Royal Park 3 | TLT | 28.3 | 1.9 | 24.0 | 31.0 |
| Royal Park 2 | TLT | 27.3 | 2.7 | 22.0 | 32.0 |
| Princes Park 2 | TLT | 21.3 | 2.2 | 17.0 | 25.0 |
| University Square 2 | TLT | 19.0 | 1.6 | 15.0 | 21.0 |
| University Square 1 | TLT | 18.3 | 1.4 | 15.0 | 20.0 |
| Lincoln Square 1 | TLT | 17.1 | 1.1 | 14.0 | 18.0 |
| High intensity mowing | TLB | 16.4 | 0.7 | 15.2 | 17.8 |
| Fitzroy-Treasury Gardens 3 | TLT | 16.2 | 1.0 | 14.0 | 17.0 |
| Fitzroy-Treasury Gardens 1 | TLT | 13.9 | 1.2 | 11.0 | 15.0 |
| Garrard Street Reserve 1 | TLT | 13.8 | 0.5 | 12.0 | 14.0 |
| Women's Peace Gardens 1 | TLT | 13.2 | 1.1 | 11.0 | 14.0 |
| Princes Park 3 | TLT | 12.2 | 1.5 | 9.0 | 15.0 |
| Murchinson Square 2 | TLT | 10.7 | 0.5 | 9.0 | 11.0 |
| Gardiner Reserve 1 | TLT | 10.7 | 1.3 | 8.0 | 13.0 |
| Pleasance Gardens 1 | TLT | 10.6 | 0.6 | 9.0 | 11.0 |
| Westgate Park 3 | TLT | 10.5 | 1.4 | 8.0 | 13.0 |
| Carlton Gardens South 2 | TLT | 10.2 | 0.9 | 8.0 | 11.0 |
| Murchinson Square 1 | TLT | 10.1 | 0.9 | 8.0 | 11.0 |
| Princes Park 1 | TLT | 9.6 | 1.2 | 7.0 | 12.0 |
| Lincoln Square 3 | TLT | 8.8 | 1.2 | 7.0 | 11.0 |
| Pleasance Gardens 3 | TLT | 8.8 | 1.2 | 6.0 | 11.0 |
| Garrard Street Reserve 3 | TLT | 7.9 | 0.4 | 7.0 | 8.0 |
| Murchinson Square 3 | TLT | 7.7 | 0.6 | 6.0 | 8.0 |
| Pleasance Gardens 2 | TLT | 7.0 | 1.0 | 5.0 | 9.0 |
| Westgate Park 1 | TLT | 4.8 | 0.8 | 4.0 | 6.0 |
| Argyle Square 3 | TLT | 4.6 | 0.6 | 3.0 | 5.0 |
| State Library of Victoria 1 | TLT | 3.9 | 0.7 | 3.0 | 5.0 |
| Westgate Park 2 | TLT | 3.8 | 0.8 | 3.0 | 5.0 |
| Women's Peace Gardens 2 | TLT | 2.9 | 0.4 | 2.0 | 3.0 |
| Lincoln Square 2 | TLT | 2.5 | 0.5 | 2.0 | 3.0 |
| Argyle Square 2 | TLT | 2.4 | 0.5 | 2.0 | 3.0 |
| University Square 3 | TLT | 2.3 | 0.5 | 2.0 | 3.0 |
| Gardiner Reserve 2 | TLT | 1.5 | 0.5 | 1.0 | 2.0 |
| Garrard Street Reserve 2 | TLT | 1.4 | 0.5 | 1.0 | 2.0 |
| State Library of Victoria 2 | TLT | 1.3 | 0.5 | 1.0 | 2.0 |
| Canning/Neill Street Reserve 3 | TLT | 1.3 | 0.5 | 1.0 | 2.0 |
| Argyle Square 1 | TLT | 0.9 | 0.3 | 0.0 | 1.0 |
| Carlton Gardens South 1 | TLT | 0.8 | 0.4 | 0.0 | 1.0 |
| State Library of Victoria 3 | TLT | 0.8 | 0.4 | 0.0 | 1.0 |
| Canning/Neill Street Reserve 1 | TLT | 0.5 | 0.5 | 0.0 | 1.0 |
| Canning/Neill Street Reserve 2 | TLT | 0.1 | 0.3 | 0.0 | 1.0 |
